# Biophysically Interpretable Inference of Cell Types from Multimodal Sequencing Data

**DOI:** 10.1101/2023.09.17.558131

**Authors:** Tara Chari, Gennady Gorin, Lior Pachter

**Affiliations:** Division of Biology and Biological Engineering, California Institute of Technology, Pasadena, California; Department of Computing and Mathematical Sciences, California Institute of Technology, Pasadena, California

## Abstract

Multimodal, single-cell genomics technologies enable simultaneous capture of multiple facets of DNA and RNA processing in the cell. This creates opportunities for transcriptome-wide, mechanistic studies of cellular processing in heterogeneous cell types, with applications ranging from inferring kinetic differences between cells, to the role of stochasticity in driving heterogeneity. However, current methods for determining cell types or ‘clusters’ present in multimodal data often rely on ad hoc or independent treatment of modalities, and assumptions ignoring inherent properties of the count data. To enable interpretable and consistent cell cluster determination from multimodal data, we present meK-Means (mechanistic K-Means) which integrates modalities and learns underlying, shared biophysical states through a unifying model of transcription. In particular, we demonstrate how meK-Means can be used to cluster cells from unspliced and spliced mRNA count modalities. By utilizing the causal, physical relationships underlying these modalities, we identify shared transcriptional kinetics across cells, which induce the observed gene expression profiles, and provide an alternative definition for ‘clusters’ through the governing parameters of cellular processes.

## Introduction

Single-cell genomics technologies are enabling high resolution, systems biology studies of cell types and their functional heterogeneity, across tissues, organs, and whole organisms [1–3]. To build a more detailed picture of how underlying DNA and RNA processing leads to cellular function and development, much focus has been given to extending these platforms to capture multiple, simultaneous measurements or ‘modalities’ at a genome-wide scale, from chromatin state information to mRNA expression and protein production [4, 5]. Such experimental approaches also provide opportunities for large-scale, mechanistic studies, to model the kinetics of these processes [6] and the roles of biological stochasticity (or noise) in driving cellular heterogeneity or differentiation [7, 8] and disease progression [9–11]. Previously, such mechanistic studies were often limited to exploring cellular processes for only a handful of genes and/or for homogeneous systems [12–14]. Thus there is an opportunity and need for using multimodal, single-cell data to gain biophysical and mechanistic insight into heterogeneous cell types [8, 15, 16].

Determination of cell types is a central task in single-cell genomics analysis [17, 18], though the definition of ‘type’ and whether designations should be of a discrete or continuous nature is a matter of debate [19, 20], and is often investigation-dependent. Here we focus on discrete categorizations, or clusters, of cells where common clustering methods include the Louvain [21] or Leiden [22] community detection algorithms (i.e. neighborhood graph-based algorithms), hierarchical clustering techniques [23], (finite-dimensional) mixture model approaches [24, 25], and marker gene-based analyses [26]. While these techniques are widely used, they are subject to heuristic tuning of hyper-parameters [17], and/or assume Gaussian-distributed data in contrast with the sparse and discrete nature of single-cell data. Clustering results are also often derived after initial dimension reduction, and assessed after further dimension reductions and embedding to 2D [27, 28]. As more modalities are introduced, delineating clusters with these methods becomes convoluted, particularly regarding the treatment of modality-specific features and variability as shown with a testicular germ cell (E13) single-cell RNA-sequencing (scRNAseq) dataset [29] (Fig. 1).

**Figure 1:**
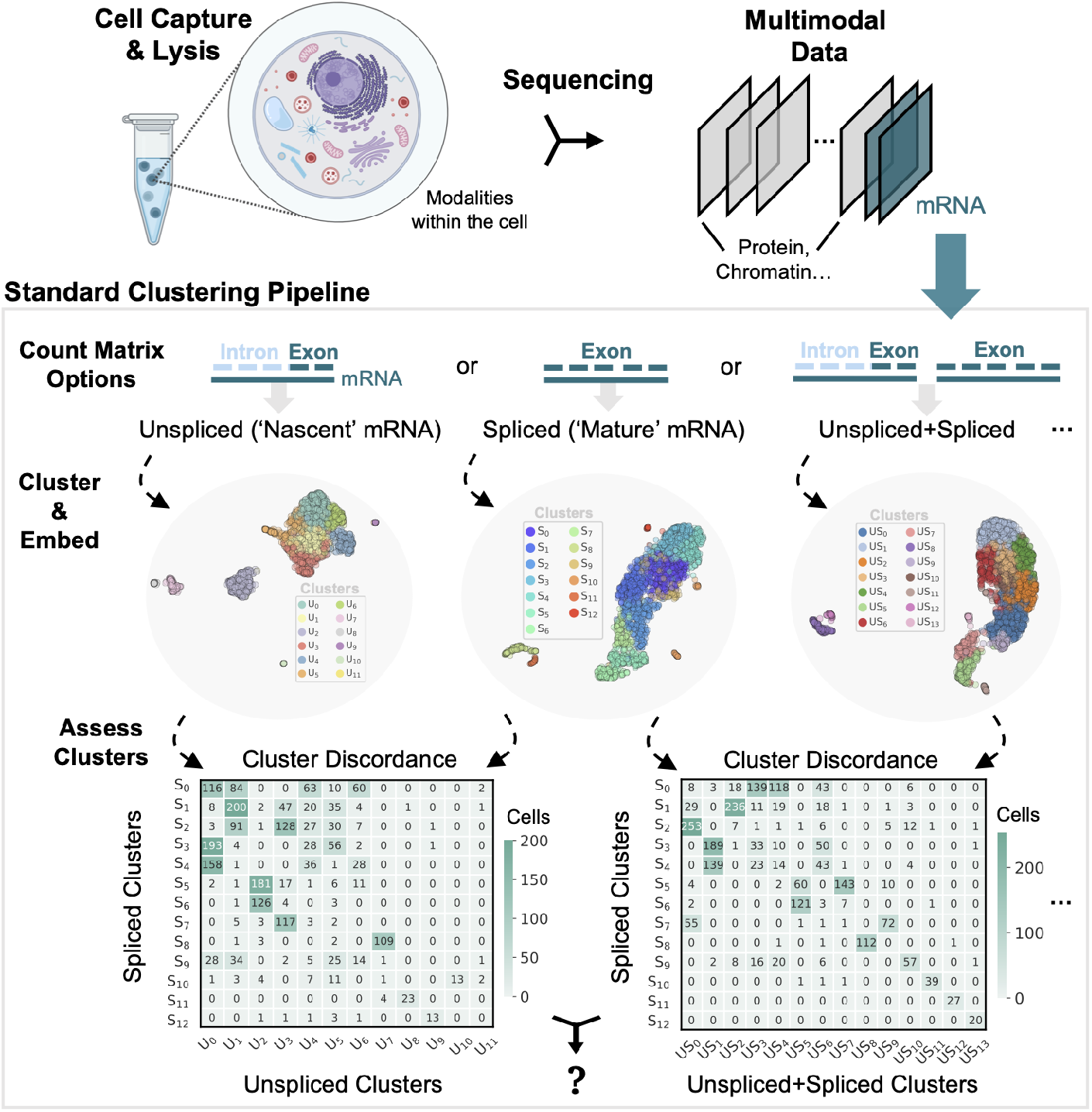
Standard Clustering Results Across Possible Count Matrix Inputs. Count matrices derived from multimodal, single-cell data with multiple mRNA count matrices (Unspliced and Spliced counts), from testicular germ cells (E13) [29]. Several matrices can be used as analysis input e.g. Unspliced counts, Spliced counts, and Unspliced+Spliced summed counts. UMAP [27] embeddings of each matrix shown, colored by Leiden [22] clusters generated independently for each respective matrix. ‘Cluster Discordance’ matrices (or confusion matrices) shown for clusters generated from Spliced counts, versus Unspliced or Unspliced+Spliced counts. Values denote counts of cells which overlap between results. Created with BioRender.com.

Specifically, for scRNAseq data, standard clustering is performed on a gene count matrix constructed from spliced gene expression, or non-intron aligning reads (representing ‘mature’ mRNA) [30]. However, two modalities, Unspliced (**U**, intron-aligning) and Spliced (**S**) molecule counts, can be obtained from most scRNAseq datasets; these are summed to generate the “count matrix” in the current 10x Genomics Cell Ranger 7.1.0 pipeline. This immediately raises the question of which matrices are relevant for clustering? The choice of matrix (Fig. 1 ‘Count Matrix Options’) has many implications for clustering methods, starting with the set of genes to be used (Supplementary Fig. 1). Perhaps unsurprisingly, different matrix choices result in distinct cluster assignments for the cells (see Methods, Fig. 1 ‘Assess Clusters’), necessitating either an arbitrary selection of one matrix to use or determination of consensus clusters, defined through some metric or heuristic, across modalities and input options.

Existing methods for determining such consensus or shared clusters largely ignore **U** counts, focusing on integrated clustering across mRNA (**S** counts), protein, and chromatin accessibility modalities [31–33]. Methods that do integrate **U** counts build on standard RNA velocity pipelines [34], which rely on large numbers of arbitrary hyperparameters and ad hoc processing steps [35], and are often incompatible with known biophysics [35]. Outside of these approaches, multimodal clustering methods often utilize heuristics to weight or balance the influence of modality-specific neighborhood-graphs or similarity matrices [33]. Alternatively, deep learning and/or embedding approaches seek to find a common space which produces the partitions of the data into clusters [31], or which can then be clustered by other existing methods [32]. Though here it is common for such methods to model the count data using discrete distributions, these methods treat the modalities as independent distributions which ignores the inherent causal relationships induced by the underlying, transcriptional processes. Furthermore, it is unclear how to justify the balance of modalities, and retain their biological variation, in such latent spaces [36, 37] (Supplementary Fig. 1). These concerns are exacerbated by additional distortions induced by upstream data transformations/reductions [38, 39].

In light of these limitations, we propose meK-Means (mechanistic K-Means) as a method to cluster cells from multimodal single-cell data under a self-consistent, biophysical model. We demonstrate meK-Means on two modalities: **U** and **S** gene count data, for which we utilize the Chemical Master Equation (CME) to formalize transcriptional processes in the cell and their governing rates [40, 41], as well as technical processes [42]. The CME provides a natural framework to model the joint distribution of **U** and **S** counts at steady-state [41]. meK-Means presents a mixture model representation of this joint distribution, to learn clusters **Z** underlying the observed gene count data (see Methods). A cluster is then inherently defined by the governing parameters of the cellular processes of interest, and represents shared transcriptional programs between genes, allowing for greater representation of gene-gene correlations as opposed to the standard, independent treatment of genes under the CME approach [43].

Thus meK-Means provides a basis for interpretable integration of modalities to learn cell clusters, and inherently balances the contribution of each modality, as well as technical and biological noise, by unifying them through their underlying, biophysical relationships.

## Results

Given a set of cell-by-gene count matrices (Fig. 2a ‘Input’), meK-Means learns clusters of cells which demonstrate shared transcriptional kinetics across genes of interest (see Methods). Specifically, given Unspliced (**U**) and Spliced (**S**) count matrices, meK-Means infers cluster-specific biophysical parameters which describe transcriptional bursting and rates of mRNA splicing and degradation (Fig. 2a ‘meK-Means Inference’), alongside learning the partitions of cells into clusters as distinguished by the parameters (see Methods, Supplementary Note). This provides an “integrated” approach to clustering with the two modalities under a CME-based, biophysical model of their causal relationships. As described in Methods, meK-Means iteratively assigns cells to a cluster, then infers parameters which induce a joint distribution of **U, S** counts recapitulating the observed counts (Fig. 2a ‘Learn Clusters’). For more discussion on the properties of the meK-Means method in relation to existing clustering [44] and mixture model approaches see the Supplementary Note.

**Figure 2:**
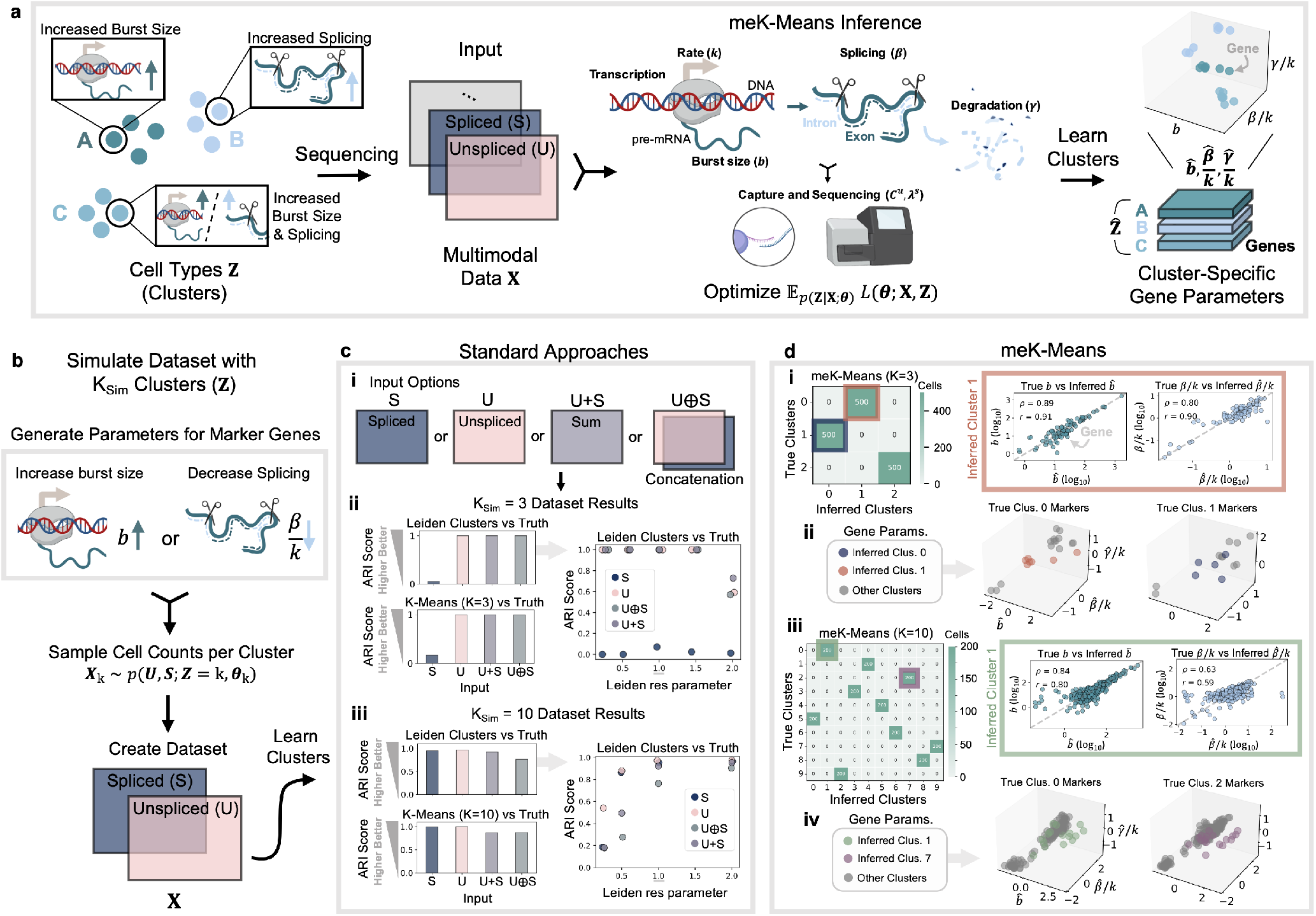
meK-Means Inference and Simulation Performance. **a)** Diagram of data generation and meK-Means inference pipeline. Input: Multimodal sequencing data with underlying clusters **Z** . Output: Inferred clusters 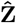with cluster-specific biophysical parameters. meK-Means model defines clusters with governing rates describing transcription and sequencing processes. **b)** Marker genes are defined for each of the K_Sim_ clusters, through either increased burst size *b* or decreased splicing *β/k* parameters (see Methods). Cell counts are sampled for each cluster and concatenated forming the dataset **X. c)** Results from standard approaches to learning cell clusters. (i) Possible input matrix options for these approaches. (ii) ARI (Adjusted Rand Index) score of default Leiden (res = 1.0) and K-Means clusters compared to the simulated truth, for K_Sim_ = 3 simulated data. Right plot shows ARI scores over different res (for Leiden) hyperparameter values. (iii) Same plots as ii for the K_Sim_ = 10 simulation data. **d)** (i) Confusion matrix showing overlap between true and meK-Means inferred clusters for the K_Sim_ = 3 simulated data. Right panel shows Spearman (*ρ*) and Pearson (*r*) correlation of inferred burst size 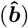 and splicing rate 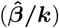 with the simulated truth. (ii) Inferred parameters for known, cluster marker genes, colored by the correspondingly marked inferred cluster. (iii) Same confusion matrix as in i for the K_Sim_ = 10 simulated data. (iv) Same parameter-space plots as in ii for the K_Sim_ = 10 simulated data. All parameter values shown in log_10_. Created with BioRender.com.

Although the number of desired clusters, K, is a parameter for the meK-Means method, meK-Means can converge to fewer than K clusters (see Methods, Supplementary Note). The parameters [*b, β/k, γ/k*] to be inferred are, respectively, transcription burst size, relative splicing rate, and relative degradation rate (relative to the rate of transcription *k*). The meK-Means model also incorporates parameters which represent technical sampling of the mRNA molecules during the sequencing process (Fig. 2a ‘Capture and Sequencing’), which are determined prior to biophysical parameter inference (see Methods).

To assess the meK-Means assignment of cells to clusters, we compared its results to the output of two popular clustering methods, Leiden [22, 45] and K-Means [46], on simulated and real single-cell data sets. Since there is no consensus on the treatment of these matrices as input for these methods, we assessed the accuracy and similarity of Leiden or K-Means results with various possible input matrices: **U, S, U** + **S, U** *⊕* **S. U** + **S** represents the summation of the individual **U** and **S** matrices, mimicking how these counts are summed in the 10x Genomics Cell Ranger pipeline 7.1.0. **U** *⊕* **S** represents the concatenation of the **U** and **S** matrices, akin to the independent utilization of modalities in many integration methods [47]. Results from these algorithms are denoted under ‘Standard Approaches’.

After obtaining clusters from meK-Means inference on real data, we extracted genes that displayed large fold-changes (FCs) in parameters (*b, β/k, γ/k*) between clusters, denoted as ‘DE-*θ*’ or ‘differentially expressed in parameter(s) *θ*’ genes. Here, DE-*θ* signifies a log_2_FC (*>* 2) in at least one parameter between two clusters (see Methods). Since the parameters describe the full, joint distribution of counts, parameter-level FCs between clusters may not be discernible FCs (log_2_FC *>* 1) at the level of mean spliced expression, which is the standard approach for differential expression [48]. Thus, DE-*θ* genes may not be DE-*µ*_*s*_, ‘differentially expressed in mean, spliced counts *µ*_*s*_’.

DE-*θ* genes were then categorized as a particular cluster’s ‘marker’ depending on which parameters were affected, though this definition is broader and does not perfectly agree with the definition of marker gene as increased, spliced gene expression. We denote a DE-*θ* gene as a cluster’s marker when burst size is increased or both splicing and degradation are decreased (suggesting increased burst frequency i.e. transcription rate *k*), as both suggest increased mRNA expression. Otherwise, an increase in splicing or decrease in degradation (increased mRNA ‘stability’) denote a marker. However, such parameter-level definitions of marker genes are flexible and likely to be task- or investigation-dependent.

### Recapitulation of Ground Truth Clusters from Simulated Datasets

To test the performance of meK-Means on datasets with ground-truth parameters and clusters, we used the model of transcription described in Fig. 2a and Methods to simulate K_Sim_ clusters, where each marker gene for the K_Sim_ clusters was distinguished by either increased burst size or decreased (relative) splicing rate (see Methods) (Fig. 2b). These gene parameters for each cluster define a probability distribution over unspliced (*U*) and spliced (*S*) molecule counts from which a dataset **X** can be sampled (see Equation (1)), containing **U** and **S** cell-by-gene count matrices.

The standard approaches (Leiden and K-Means) were then run with the simulated data **X** as input (Fig. 2c i). To assess the resulting cluster assignments, the Adjusted Rand Index (ARI) score was used to measure the similarity between the approach’s assignments and the true clusters from simulation, where 1.0 denotes overlapping assignments and 0.0 represents poor or random assignments. For the case where K_Sim_ = 3, both Leiden and K-Means could not determine the true clusters with **S** counts only (ARI score near 0.0), though the assignments were largely correct upon the inclusion of **U** counts (i.e. most of the information is in the **U** counts) (Fig. 2c ii). For another K_Sim_ = 3 simulation, where only burst size was increased for marker genes, we found that all input options except **U** counts only could discover the original clusters (Supplementary Fig. 2a). However, as the resolution (‘res’) hyperparameter for Leiden was increased, cluster assignment for these simulated datasets became increasingly inaccurate across all input matrices (Fig. 2c ii, Supplementary Fig. 2a). At low res values, the **U** and **S** matrices also displayed poor cluster assignment (Fig. 2c ii, Supplementary Fig. 2a).

In the K_Sim_ = 10 case, we found that the input matrices with the most accurate clustering results were the individual count matrices **U** and **S**, with diminished accuracy for **U** + **S** and **U** *⊕* **S** (Fig. 2c iii). Additionally, larger res values resulted in greater similarity to the ground truth clusters, in contrast to the K_Sim_ = 3 simulations (Fig. 2c iii). We also tested these approaches on a negative control simulation, where K_Sim_ = 1 and thus all cell counts are sampled from the same set of gene parameters. Across res values for Leiden, or K values for K-Means, most cluster assignments did not converge to the single cluster, and the assignments differed dramatically between the input options even at the same res or K value (Supplementary Fig. 3a). Together, this demonstrates instabilities in these approaches, both across hyperparameter values and input matrices, and opaque sensitivities to dataset-specific features, given simulated data with underlying cell type or cell cluster structure.

In contrast, meK-Means was able to recover the underlying clusters for the simulated cases, converging on the correct number of clusters even over a range of Ks, inherently utilizing both **U** and **S** matrices as input (Fig. 2d i,iii, Supplementary Fig. 2b). This was additionally the case for the K_Sim_ = 1 simulation, where various values of K converged to K=1 cluster (Supplementary Fig. 3b). Along with the correct clusters, the learned physical parameters were highly correlated with the ground truth parameters (Fig. 2d i,iii). As shown in Fig. 2d ii and iv, for the K_Sim_ = 3 and K_Sim_ = 10 simulations respectively, the inferred parameters for the known marker genes clearly distinguish the ‘marked’ cluster from the other clusters’ parameters (in grey), as expected. This demonstrates, for these well-defined test cases, recovery of both the correct partitions or clusters of cells and the kinetic parameters themselves.

### Kinetic Analysis of Motor Cortex Cell Types

To assess meK-Means results on biological data, we first utilized the Allen Institute Mouse Primary Motor Cortex (MOp) dataset [23]. This contained highly granular cell annotations produced from hierarchical, iterative clustering on the spliced expression matrix [23]. meK-Means was run on a subset of 4 cell types which displayed a range of population sizes (from 80 to 1,300 cells), across 1,000 highly variable genes (HVGs) selected with scanpy (see Methods) (Fig. 3a).

**Figure 3:**
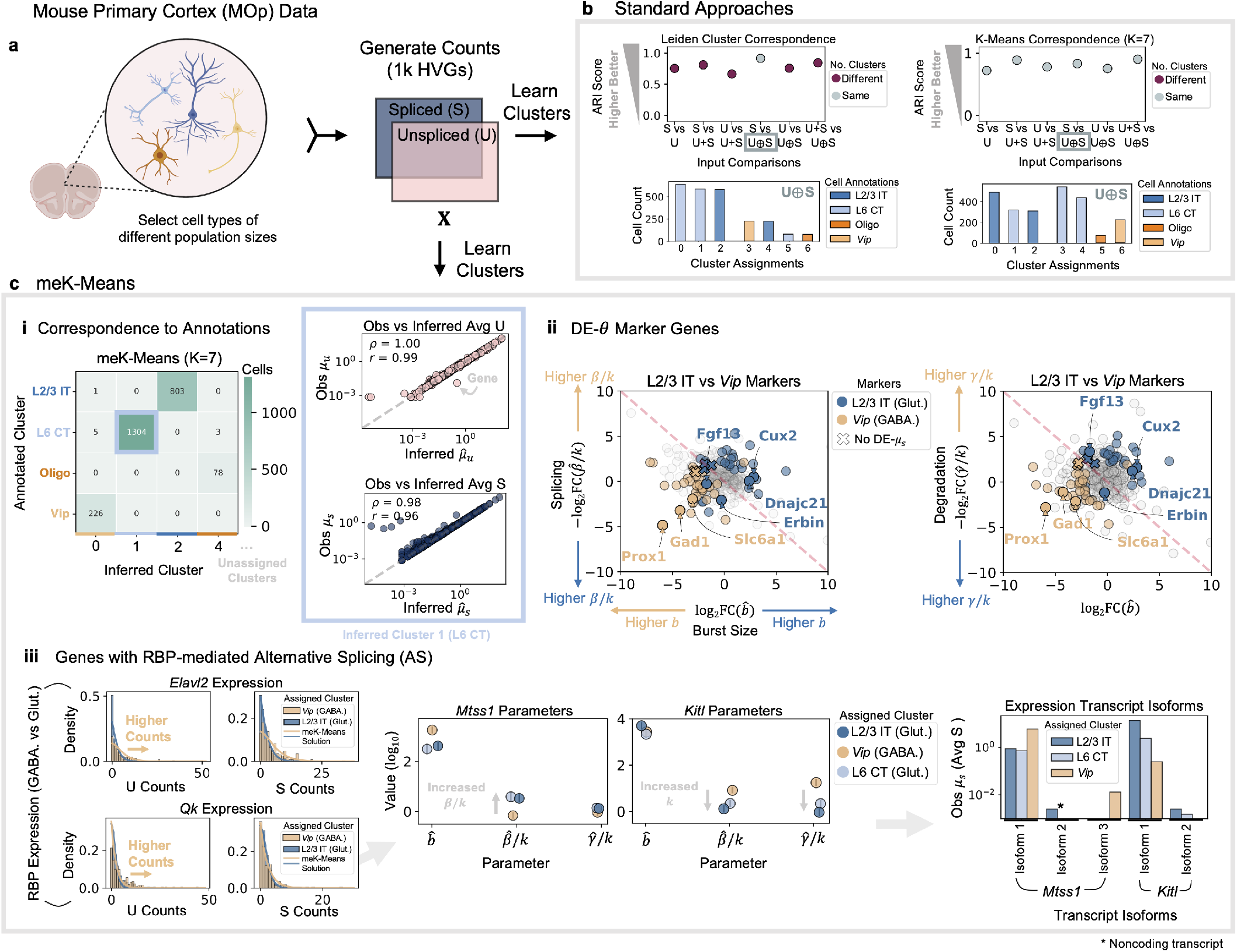
meK-Means Inference for MOp Data. **a)** Cell types of various population sizes selected from the Mouse Primary Cortex (MOp) dataset [23]. **U**,**S** count matrices generated from the data. Gene counts from the top 1,000 highly variable genes (HVGs) used (see Methods). **b)** Results from standard approaches to learning clusters. Left plots show ARI scores between each Leiden result, across the possible input options. Result pairs with differing numbers of clusters denoted. Distribution of original annotations across assigned clusters shown for the **U** *⊕* **S** input. Right plots show same results for K-Means with K=7, matching the number of Leiden clusters. **c)** (i) Confusion matrix showing overlap between original annotations and inferred clusters from meK-Means with K=7. Only 4 clusters assigned. Right panel shows Spearman (*ρ*) and Pearson (*r*) correlation of the observed **U**,**S** means (e.g. *µ*_*u*_) versus inferred means 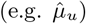. (ii) ‘DE-*θ*’ or ‘Differentially-Expressed in parameter(s) *θ*’ genes, based on log_2_ fold-change (FC) between clusters (see Methods). Left plot shows splicing vs burst size FCs, and right shows degradation vs burst size FCs of L2/3 IT versus *Vip* neurons (assigned from inferred clusters). Genes denoted with Xs in the plot indicate DE-*θ* genes where FCs were not detected at the (observed) mean spliced-level (i.e. not DE-*µ*_*s*_). (iii) (Left to Right) Raw **U**,**S** count histograms for two RBPs (RNA-Binding Proteins) with analytical solution curves from meK-Means parameters overlaid, in a GABAergic (GABA.) cluster, *Vip*, and a Glutamatergic (Glut.) cluster, L2/3 IT. Inferred parameters for *Qk* (RBP)-mediated genes in GABA. and Glut. clusters, error bars indicate 99% C.I. (see Methods). Transcript isoform expression for the *Qk*-mediated genes. Created with BioRender.com.

For the results of the standard approaches, we calculated ARI scores between cluster assignments generated from the pairs of possible input options. These comparisons evaluate the stability and consistency between the generated clusters, mimicking how with real datasets we often do not have ‘ground truth’ and use such algorithms to decide what partitioning of the data to retain. This is akin to how ‘consensus’ clusters are selected i.e. as clusters that appear the most concordant between modalities, where concordance is assumed to represent stable cluster structure [39]. Leiden results, at the default resolution, yielded 7 clusters, for the most consistent pair of input options **S** and **U** *⊕* **S** (Fig. 3b). K-means at the same resolution, K=7, resulted in different distributions of the cells and original cell annotations across the clusters, where **U** + **S** and **U** *⊕* **S** were the most similar inputs (Fig. 3b). Input option pairs also often converged on different numbers of clusters, and larger res hyperparameter values caused merging of clusters and generation of new clusters with no designated annotations in the original study (even in comparison to the existing, sub-type annotations) (Supplementary Fig. 4a). As described with the simulated datasets, this suggests instability in the cluster determinations, counter-productive for determining consensus clusters across input options. A user would then require prior-knowledge of the expected clusters to select the ‘best’ resolution hyperparameter, input matrix, or both.

We then ran meK-Means, with K=7, and converged on 4 clusters corresponding to the original cell annotations: L2/3 IT, L6 CT and *Vip* neurons, and Oligodendrocyes (Oligo) (Fig. 3c i). Convergence to these four clusters was also stable over several values of K, and displayed a better Akaike Information Criterion (AIC) value, measuring model quality through information loss, versus smaller numbers of clusters (Supplementary Fig. 4b). This provides a metric based on the model’s fit to the data to determine what number of clusters are discernible for the given input. Though we do not have ground truth parameters to assess meK-Means, from the inferred physical parameters we calculated the moments (means) of each gene’s unspliced and spliced counts 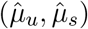 (see Equation (3)) and found they were highly correlated to the observed unspliced and spliced means, *µ*_*u*_, *µ*_*s*_ (Fig. 3c i, Supplementary Fig. 5a) (see Methods).

We then identified DE-*θ* marker genes between the inferred cell clusters. These markers aligned with markers found previously by the *Monod* package (for single-cell, CME-based inference) [37], where the same model of transcription (Fig. 2a) was fit on all GABAergic (GABA.) and all Glutamatergic (Glut.) cells, as delineated by the original cell type annotations [23]. This included increased burst size for *Dnajc21* in the Glut. cells (Fig. 3c ii), increased burst size for *Nectin1* in the GABA. cells (Fig. 3c ii), and increased splicing for *Erbin* in the Glut. cells (Fig. 3c ii, Supplementary Fig. 6 top row). The DE-*θ* markers also encompassed known markers for L2/3 IT and *Vip* neurons, such as *Cux2* and *Prox1* respectively (Fig. 3c ii) [23]. Established markers were also extracted for the other clusters, including between the two Glut. clusters (e.g. *Slc30a3* and *Cux2* for L2/3 IT neurons and *Foxp2* and *Sulf1* for L6 CT neurons) (Supplementary Fig. 6) [23]. We additionally found DE-*θ*, but not DE-*µ*_*s*_, genes (Fig. 3c ii, Supplementary Fig. 6) where for example, degradation rates were lower for the growth factor *Fgf13* in both L2/3 IT and L6 CT (Glut.) neurons compared to *Vip* (GABA.) neurons (Fig. 3c ii), but such differences were not necessarily detectable between *µ*_*s*_ (mean spliced expression) values (Supplementary Fig. 6 top row). Distinguishing clusters of cells by differing degradation rates is also particularly important for understanding disease development, as has been described with Alzheimer’s-related alterations of mRNA decay rates and stability in affected cells [9].

Alternative splicing (AS) regulation in the brain is another area of interest for mechanistic studies, as AS contributes to determining cellular identity and connectivity between cells [49, 50], and differential AS has been studied between GABA. vs Glut. populations [50]. With the meK-Means results, we can uncover interesting patterns in the behavior of genes whose AS is mediated by certain RNA Binding Proteins (RBPs) such as *Elavl2* and *Qk*, which tend to be more highly expressed in GABA. populations (Fig. 3c iii). The inferred parameters for these RBPs additionally produced marginal distributions which recapitulated their observed **U**,**S** count distributions (Fig. 3c iii, Supplementary Fig. 5b).

In particular, between GABA. and Glut. cells, the RBP *Qk* can mediate *Mtss1* exon inclusion in GABA. cells and *Kitl* exon inclusion in Glut. cells [50]. From the inferred parameters for these target genes, we found increased splicing for *Mtss1* and increased burst frequency (decreased splicing and degradation) for *Kitl*, in Glut. clusters (Fig. 3c iii). Previous studies have demonstrated the regulation of burst frequency for increasing and decreasing exon skipping/inclusion in AS [51], though there is less work on the relationship between splicing rate and AS [52], suggesting an alternative avenue of investigation regarding AS regulation. The changes in splicing rate for *Mtss1* were also noted in previous GABA. versus Glut. population-level analysis with the *Monod* package [37]. These patterns of AS were also mirrored in the differential isoform expression of *Mtss1* and *Kitl* transcripts between the GABA. and Glut. clusters (Fig. 3c iii).

### Alternative Identification of Clusters in PBMC Data

To test meK-Means inference on a less structured system of cells, we used the 10x Genomics 10k Human PBMC (Peripheral Blood Mononuclear Cell) v3 dataset, which provided only rough, initial annotations of general clusters: T cells, B cells, and Monocytes (see Methods) [42]. We additionally expanded the repertoire of genes used for inference beyond HVGs, as it is not clear that the best option for multimodal data is to select genes solely based on their spliced variance. We instead selected 1,082 genes from a list of genes associated with PBMC subtypes, gathered over several studies and various experimental methodologies [53] (Fig. 4a) (see Methods).

**Figure 4:**
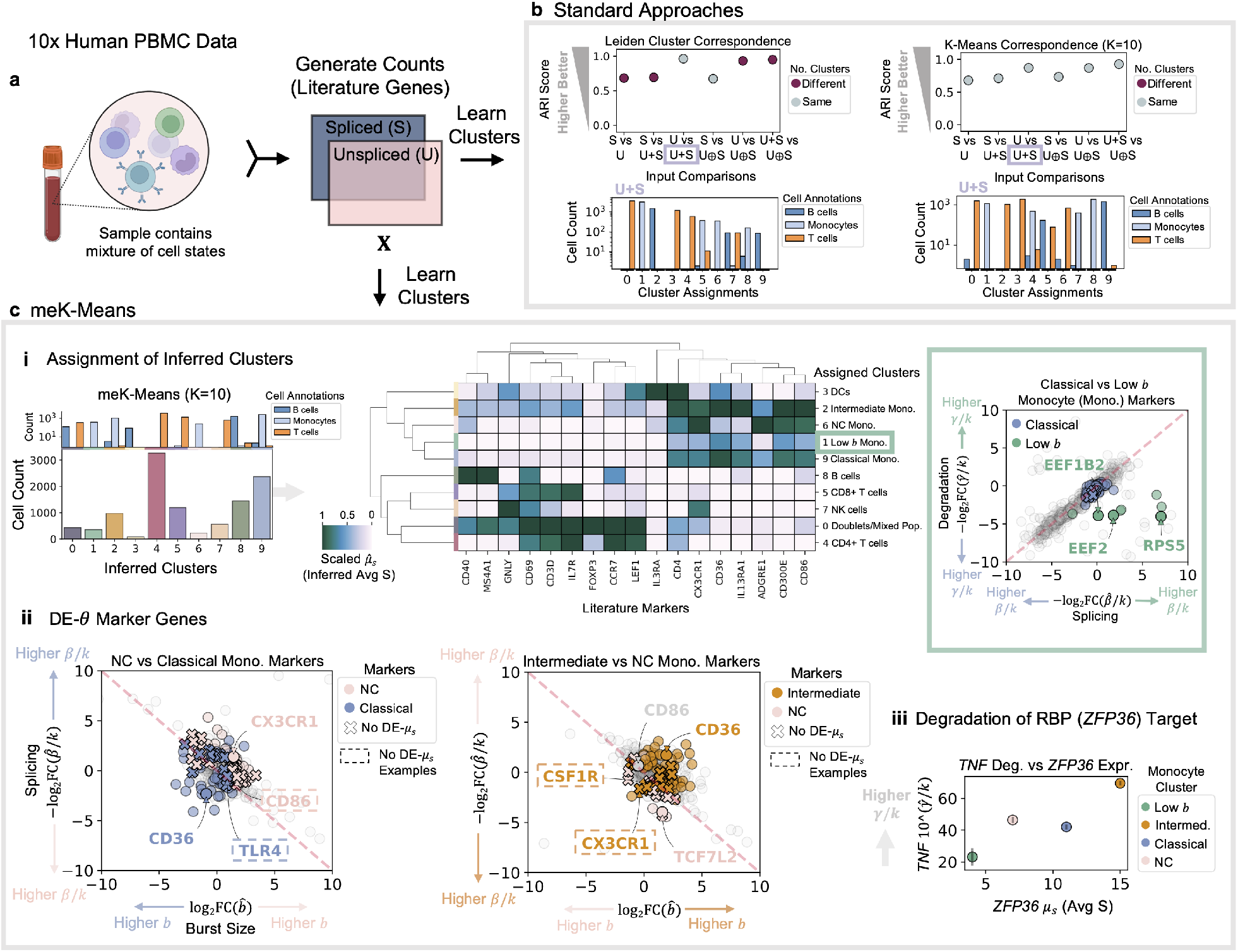
meK-Means Inference for PBMC Data. **a) U**,**S** count matrices generated from 10x Human Peripheral Blood Mononuclear Cells (PBMCs). PBMC-relevant genes from the literature used for inference (see Methods). **b)** Results from standard approaches to learning clusters. Left plots show ARI scores between Leiden result pairs, across the possible input options. Pairs with differing numbers of clusters denoted. Distribution of original annotations over learned assignments shown for the **U** + **S** input. Right plots show same results for K-Means with K=10. **c)** (i) Barplot of cell counts for meK-Means inferred clusters (K=10). Top plot shows distribution of original annotations across these clusters. Middle panel shows heatmap of the inferred clusters organized by inferred spliced mean expression 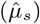 of known PBMC markers. Expression scaled for each gene/column. Right panel shows DE-*θ* marker genes between the Classical and Low *b* monocyte (Mono.) clusters (see Methods). Axes show FCs in degradation versus splicing. (ii) DE-*θ* genes between monocyte clusters (see Methods). Left plot shows splicing versus burst size FCs for assigned Non-Classical (NC) versus Classical monocytes. Right plots shows same FCs for Intermediate versus NC monocytes. Marker genes denoted with Xs were not DE-*µ*_*s*_ (example genes in dashed lines). (iii) Degradation rates for *TNF* versus mean (observed) spliced expression of *ZFP36* (RNA Binding Protein, RBP), shown across monocyte clusters. Error bars indicate 99% C.I. (see Methods). Created with BioRender.com.

Default Leiden clustering produced 10 clusters, for the most consistent pair of inputs **U** and **U** + **S**, though at the same res value, the number of clusters varied depending on the input option (Fig. 4b). K-Means at K=10 yielded clusters with different proportions of the original cell annotations across the assignments, as compared to the Leiden results for the same input, and the inputs **U** + **S** and **U** *⊕* **S** were the most similar rather than **U** and **U** + **S** (Fig. 4b).

Assignment from meK-Means for K=10 yielded 10 clusters (Fig. 4c left), which additionally showed a better AIC value across the K values tested (Supplementary Fig. 7a). For initial assessment of the clusters, we used the 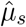mean count approximations (see Equation (3)) in each cluster to assign preliminary annotations based on gene expression of literature markers (Fig. 4c i middle). DE-*θ* genes between the inferred clusters encompassed both general marker genes (delineating T cells, B cells, and Monocytes) (Supplementary Fig. 7b) as well as established markers for populations such as Natural Killer (NK) cells and Dendritic Cells (DCs) (Supplementary Fig. 8a). We also detected a doublet/mixed population of originally labeled T and B cells, discernible by dual expression patterns of T and B cell markers, though the mixed population may reflect a limitation of genes selected for inference (Supplementary Fig. 8b).

Interestingly, despite canonical monocyte markers such as *CD14* and *CD16* being filtered out (not included) in inference due to low **U** counts, meK-Means was able to discern several monocyte clusters: a low burst size (Low *b*) population of classical monocytes, classical monocytes, intermediate monocytes, and non-classical (NC) monocytes (Fig. 4c i middle, Supplementary Fig. 7c). Low burst size monocytes (Supplementary Fig. 8c) displayed increased splicing (and decreased degradation rates) for genes related to ribosome production and protein synthesis (Fig. 4c i right), compared to classical monocytes. Between the other monocyte clusters, *CD86* was a DE-*θ* gene which distinguished NC and classical monocytes (Fig. 4c ii left), and has recently been reported as an alternative marker for distinguishing these monocytes subtypes [54]. Notably, for genes such as *CD36* and *CX3CR1*, studies have found it hard to detect differential expression between these subtypes at the level of spliced counts [54], yet they display differential behavior across monocyte clusters in their inferred parameters (even when they are not DE in *µ*_*s*_) (Fig. 4c ii). We additionally found other examples of DE-*θ* but not DE-*µ*_*s*_ gene markers between monocyte clusters, such as *TLR4* and *CSF1R* (Fig. 4c ii), which are relevant to macrophage commitment and activation, and display subtype specific behavior in various disease conditions [10, 55].

Across these monocyte clusters, we also explored relationships between expression of RBPs and their targets genes, where for example, with increasing expression of *ZFP36* we see an increase in the degradation (or decay) rates of *TNF* (Fig. 4c iii), following previous literature on the role of *ZFP36* in increasing *TNF* decay in immune cells [12, 56, 57]. In general, how to define more granular monocyte clusters beyond the broad categorizations of ‘classical’, ‘intermediate’, and ‘non-classical’, remains an open question [58]. However, meK-Means offers an alternative avenue for studying such clusters or populations, moving beyond spliced gene expression analysis to utilize the modalities available within scRNAseq data and extract differences in the data generating processes themselves, grounded by the biophysics of the system.

## Discussion

MeK-Means offers a scalable and interpretable methodology for defining clusters in single-cell, multimodal datasets through the governing parameters of their underlying, cellular processes. As these parameters describe the full, joint distribution of the data, we can identify differences between cells not discernible from standard, mean expression analysis. This enables identification of marker genes based on differential kinetic behavior, and exploration of mechanistic trends across genes and clusters.

The current workflow requires a selection of genes, from which the biophysical parameters and clusters are inferred, as well as a user-defined value of K. Metrics such as AIC can be used for selecting a model across a range of K values, or alternatively, as K upper-bounds the number of clusters learned, a high K value can be set, allowing meK-Means to converge to a possibly smaller number of clusters. The selection of genes for inference is also important, as genes without variability across cells can impede clustering. However, with multimodal data, common practice of selecting HVGs from spliced gene expression may not be the most informative approach, particularly if there is variability in another modality. Though we demonstrate an alternative approach of choosing genes from across the literature, in the future one could learn which genes to retain for discerning clusters during inference, akin to determining which genes contribute most to a particular task or objective function such as separation of cell types [59, 60].

Another limitation of the current meK-Means structure is that through our inference procedure we lose single-cell resolution at the parameter-level; each gene has one set of parameters per cluster. This highlights the importance of obtaining biological replicates for statistical analysis between clusters. By combining the meK-Means framework with previously demonstrated machine learning techniques for scalable CME inference, one could in principle, retain single-cell *and* single-gene resolution [61]. Such an approach would generalize the finite-dimensional representation of the cells in meK-Means to continuous representations of cells, where each cell is a nondeterministic function of a latent representation [61]. It would also incorporate the effects of cell-size on gene expression [61, 62], assumed to be negligible or incorporated in cluster identity under the meK-Means model.

However, meK-Means results can inherently be combined with a host of statistical tools to analyze the inferred parameters and model power. For example, the Fisher information matrix calculated from the inferred models (see Methods) can be used to assess uncertainty in parameter estimates, as well as information content of parameters, to optimize downstream experiment design [63, 64]. Additionally, Chi-squared goodness-of-fit testing can be used to reject genes with poor parameter inference (see Methods).

MeK-Means is also integrated in the *Monod* package for CME inference from single-cell data [37], where different models of transcription can be tested and their parameter fits on the data analyzed, to determine if particular models of transcription fit certain genes better than others (or if certain models cannot be distinguished) [65]. Though the meK-Means model currently only considers bursty transcription, splicing, and degradation of mRNA, such models can be extended in a self-consistent manner to include other increasingly popular readouts such as chromatin accessibility data and protein expression [13, 41]. Extension of meK-Means with machine learning techniques, as mentioned above, also enables parameter inference for CME models without exact solutions [61, 66]. Such extensions are particularly exciting given more recent developments in ‘unbiased’ sequencing technologies e.g. random primer-based [67] or long-read sequencing [68], which could broaden transcript capture and retention of transcripts in various states of processing. The meK-Means framework can also be extended to incorporate data from multiple batches, where the technical sampling parameters can be learned or set *a priori* per batch, with inferred, physical parameter values comparable across batches.

Overall, meK-Means demonstrates a ground-up approach to clustering multimodal data based on shared transcriptional kinetics. The method unifies modalities, and provides a biophysically coherent, interpretable avenue for representing single-cell data and enabling hypothesis-driven biological investigation.

## Methods

### Length-Bias, CME Model of Sequencing Data

The Chemical Master Equation (CME) formalism can be used to describe the probability distribution of molecule counts over time, given some rates of transition between states, particularly for low-count (discrete) systems [40, 41]. It has been used extensively in the fluorescence transcriptomics literature over the past couple of decades to model the processes of transcription and protein synthesis in various cell systems [13, 40, 41, 69]. Briefly, the CME defines the differential equation *dp*_*X*_*/dt* describing the time evolution of *p*(*X, t*), or the probability distribution at time *t* over the random variable(s) *X* e.g. *U, S* or Unspliced, Spliced molecule counts. We study the behavior of the probability distribution over discrete molecule counts *U*_*g*_, *S*_*g*_ for each gene *g* at time *t*, defined as *p*(*U*_*g*_, *S*_*g*_, *t*), in the long-time limit (steady state), *p*(*U*_*g*_, *S*_*g*_) as *t → ∞*.

In particular, we utilize a CME-based model of transcription developed in [42], reproduced here (see Fig. 5), which models the bursty transcription, splicing, and degradation of mRNA molecules as well as molecule sampling from the technical capture and sequencing processes, to account for length-biased capture noted in poly-A tail based sequencing techniques such as with 10x Genomics. This is the model from which simulated data is generated (see Simulation), and which defines the parameters for meK-Means inference.

**Figure 5:**
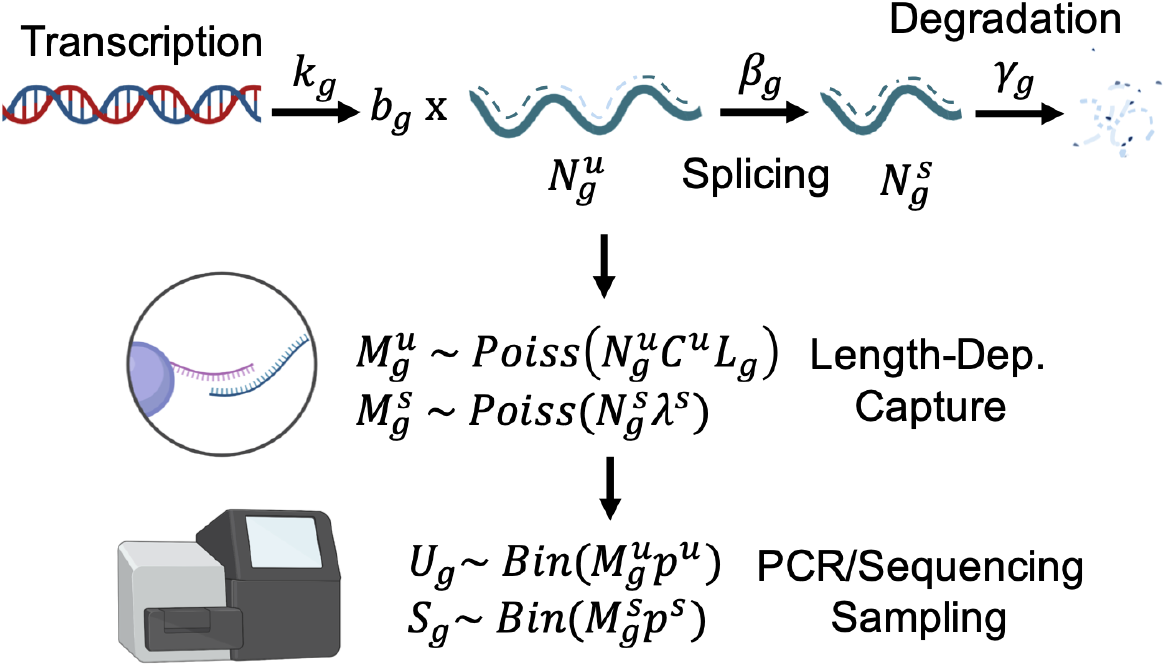
Technical, Length-Bias Model for meK-Means. The diagram shows gene-specific rates to be inferred that govern transcriptional processes, as well as the sampling processes which occur during transcript capture, e.g. during 10x Genomics poly-A based capture, and sequencing/library PCR. Created with BioRender.com.

As described in [42], certain gene parameters are not independently identifiable, thus we define relative splicing and degradation parameters, *β*_*g*_*/k*_*g*_ and *γ*_*g*_*/k*_*g*_, where splicing and degradation rates respectively are relative to the transcription rate *k*_*g*_. *b*_*g*_ represents the mean of geometrically-distributed bursts of transcription [70]. For the global (non-gene-specific) technical parameters, the *Poiss* and *Bin* capture and sequencing sampling parameters (see Fig. 5) are also not independently identifiable, thus we define net sampling rates *λ*^*u*^, *λ*^*s*^ (where *λ*^*u*^ = *C*^*u*^*L*_*g*_, *L*_*g*_ represents length of the gene) which contain *p*^*u*^, *p*^*s*^. For simulation and meK-Means inference, the global technical parameters are set prior to data generation or inference of the physical parameters (i.e. we do not perform a grid search over these parameters during meK-Means inference) (see Simulation and Dataset Preprocessing).

With this CME formulation we can define the steady-state probability generating function (PGF) form, *H*, of *p*(*U*_*g*_, *S*_*g*_), and solve for *p*(*U*_*g*_, *S*_*g*_; ***θ***_*g*_) where ***θ***_*g*_ = [*b*_*g*_, *β*_*g*_*/k*_*g*_, *γ*_*g*_*/k*_*g*_, *λ*^*u*^, *λ*^*s*^] as described in [41, 42]:

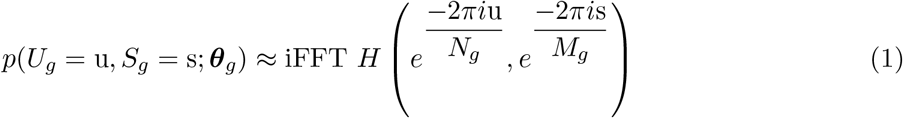

where u = 0, .., *M*_*g*_ *−* 1, s = 0, .., *N*_*g*_ *−* 1 and *N*_*g*_, *M*_*g*_ are sufficiently large, positive integers. This amounts to evaluating the PGF *H* around the complex unit circle and performing an inverse Fourier transform (denoted iFFT) to obtain the molecule count probabilities [41]. For a dataset **X**, which comprises matrices **U** and **S** *∈* ℝ^N*×*G^ (for N cells and G genes), *N*_*g*_ and *M*_*g*_ are defined as *N*_*g*_ = max(**U**_*g*_), *M*_*g*_ = max(**S**_*g*_).

To obtain MLE parameter estimates, 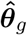 given **X**, the Kullback-Leibler Divergence (KLD) between the observed molecule count distribution and the distribution generated under the CME model is minimized :

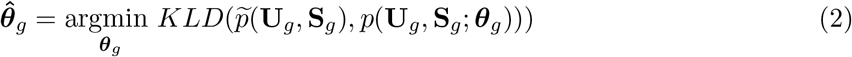

where 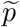 describes the observed histogram of counts. To optimize the global parameters *λ*^*u*^, *λ*^*s*^ a grid search can be performed where Equation (2) is optimized for possible pairs of (*λ*^*u*^, *λ*^*s*^), and an optimal pair of (*λ*^*u*^, *λ*^*s*^) are chosen with the minimum KLD. This grid search is performed on the datasets (see Dataset Preprocessing) prior to meK-Means inference, to obtain and set reasonable *λ*^*u*^, *λ*^*s*^ values.

The moments of the CME model (Fig. 5) are derived in [42]. For this study, we utilize the moment derivations for the unspliced and spliced mRNA count means:

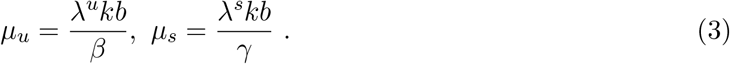

A general framework and tools for performing the numerical integration and parameter optimization for CME model inference with single-cell data are implemented in the *Monod* package [37]. meK-Means (see meK-Means Model) is integrated within the *Monod* package, utilizing these established workflows for solving such CME-based systems.

## Simulation

For all simulations, the technical parameters were set to *C*^*u*^ = *−*7.15, *λ*^*s*^ = *−*1.525 in log_10_. All physical, gene parameters were initially sampled from a multivariate Gaussian with *µ* = [1.2, 0.2, 0.5, 5], *σ*= [0.6, 0.3, 0.5, 0.5] representing [*b, β/k, γ/k, L*], in log_10_. Covariances between *b, β/k, γ/k* were 0.8, and a slightly negative covariance of -0.1 was set between *b* and *L* (mimicking the findings in [16]).

For the K_Sim_ = 3 simulation, parameters for 100 genes were sampled, with 10 unique marker genes per cluster k. For each marker gene, noise *ϵ ∼ Normal*(1.0, 0.1) was added to *b* with probability = 0.1, otherwise *ϵ* was subtracted from *β/k*. We additionally simulated the case where only *b* was altered for marker genes. 500 cells (observations) were then sampled for each cluster k, from the distribution of molecules counts defined as in Equation (1).

For the K_Sim_ = 10 simulation, parameters for 500 genes were sampled, with 20 unique marker genes per cluster k. For each marker gene, noise *ϵ ∼ Normal*(1.0, 0.1) was added to *b* with probability = 0.2, otherwise *ϵ* was subtracted from *β/k*. 200 cells (observations) were then sampled for each cluster k, from the distribution of molecules counts defined as in Equation (1).

Thus we simulate marker genes with increased ‘expression’ i.e. increased burst size or decreased splicing. We additionally chose the perturbation of splicing rates to denote markers, as the *U, S* modalities are particularly suited to studying changes in splicing.

### Dataset Preprocessing

For the demonstration of results from standard clustering pipelines (Fig. 1), the unspliced (intron-aligning) and spliced (non-intron-aligning) mRNA count matrices were taken directly from the original study [29] and concatenated into a single loom file for processing. To mimic standard processing steps, the rows or cells of each input matrix was normalized to a total of 10^4^ counts and log1p-transformed. 2000 HVGs were selected for each matrix with scanpy [45] highly variable genes, and each matrix reduced to 40 dimensions with PCA prior to clustering. UMAP [22] was run on these reduced matrices at default settings to produce the 2D embeddings.

Datasets selected for meK-Means analysis were previously described and processed from the original FASTQ files in [61] and [42]. For both the Allen Institute Mouse Primary Motor Cortex (MOp) dataset [23] (processed in [61]) and 10x v3 10k Human Peripheral Blood Mononuclear Cell (PBMC) dataset (https://www.10xgenomics.com/resources/datasets/10-k-pbm-cs-from-a-healthy-donor-v-3-chemistry-3-standard-3-0-0) (processed in [37]), count matrices were generated with *kallisto*|*bustools* 0.26.0 [71, 72]. Specifically, the kb ref command with the --lamanno option was used to build exonic and intronic references for each genome. kb count with the --lamanno workflow and the -x 10xv3 technology option was then used to obtain unspliced and spliced mRNA count matrices [42, 61]. These raw count matrices were then used as input for meK-Means. The mm10 and GRCh38 (2020-A version) reference genomes were used for pseudoalignment, downloaded from 10x Genomics (https://support.10xgenomics.com/single-cell-gene-expression/software/downloads/latest). The datasets were also selected as they displayed either high-resolution cell annotations or rough, initial assignments only.

These two datasets were first filtered for cell barcodes with at least 1,000 (MOp data) or 3,000 (PBMC data) molecular barcodes total (across genes) prior to gene filtering. The MOp dataset was additionally filtered for four cell types, which displayed different population sizes, incorporating two Glutamatergic, one GABAergic, and one non-neuronal population. 2,000 HVGs were then selected with scanpy [45] highly variable genes [61]. meK-Means inference was then run on a filtered subset of 1,000 HVGs. Genes were filtered out based on a minimum and maximum number of unspliced or spliced, **U** or **S**, counts: *µ*_*u*_, *µ*_*s*_ *≤* 0.01, max(**U**), max(**S**) *≤* 3, or max(**U**), max(**S**) *≥* 400 (as described in [37]). For the PBMC dataset, a gene list of 2,357 PBMC-related markers and genes of interest (from across the literature) was used [53]. 1,082 genes remained after the same filtering procedure was applied, and were used for meK-Means inference. All dataset download links are provided in Supplementary Table 1.

To determine the global, technical parameters for each dataset prior to meK-Means inference (see ‘Capture and Sequencing’ in Fig. 2a), a grid search over pairs of technical parameters was run (as outlined in [37]). For each grid point or pair of unspliced/spliced sampling parameters, an optimization of the biophysical parameters was performed (see Equation (2)). This grid search was run independently for subsets of cells in different cell types (from existing annotations). An ‘optimal’ pair of technical parameters was selected for each cell subset i.e. the pair under which the MLE (biophysical) parameter estimates had the minimum KLD summed across genes (see Equation (2)). The centroid or ‘consensus’ pair of technical parameters for the whole dataset was then determined from across the subset-specific, optimal parameter pairs. This approach to technical parameter estimation could also be performed across random subsamples of cells (across different cell types), using housekeeping or non-variable genes.

### meK-Means Model

The meK-Means model introduces the latent variable *Z* to the CME model of transcription above, expanding the likelihood model of the data from *p*(*U, S*|***θ***) to *p*(*U, S, Z*|***θ***) . Here *Z* can take any (integer) value from 1 to the user-defined K. Given that both the posterior, *p*(*Z*|*U, S*, ***θ***), and parameters, ***θ***, are unknown, we take an Expectation Maximization (EM)-based approach to optimize the Q function:

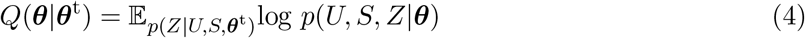

iterating between updating the posterior given parameter estimates ***θ***^t^ (E-step), and determining the MLE parameter estimates which then maximize *Q*(***θ***|***θ***^t^) (M-step). See Algorithm 1 below. The multi-dimensional nature of the input data and the non-independent treatment of the dimensions (modalities) also naturally contextualizes the meK-Means EM approach as a joint clustering method of high-dimensional, tensor data [25] (see Supplementary Note for further discussion).

To initialize the posterior, *p*(**Z**|**U, S, *θ***), given count matrices **U, S**, K-Means clustering was performed on the **S** matrix (for the user-defined K) and for each cell n, 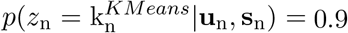 where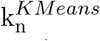 is the cluster k assigned to cell n by K-Means. For all other k, 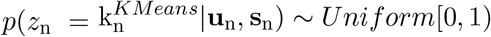

Since the numerical procedure for obtaining parameter estimates for the defined CME model requires a histogram over observed counts (see Equation (2)), we use hard assignment of cells to a single latent state or cluster k during the M-step, where the cell is assigned k such that

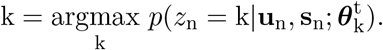

This is akin to the hard assignment of each observation (cell) to a cluster centroid (based on distance from the observation to that centroid) in K-Means clustering (see Supplementary Note for comparison to K-Means).

Note that in Equation (7), given that the hard assignment of k is determined by the maximum posterior value for a given cell, meK-Means can converge to a final number of assigned clusters less than the upper bound K set by a user.

All approaches, i.e. Leiden, K-Means, and meK-Means, were run on an Intel Xeon Gold 6152 at 2.10 GHz using up to 30 cores. Inference for the *∼*1,000 genes was run in parallel, for a compute time of 1 min per epoch. An example Google Colaboratory notebook for meK-Means analysis is also available in the Github repository, which can be run for free in the Google Cloud.

### Standard Approaches for Clustering

We compare the results of meK-Means to the outputs of two popular methods in single-cell data analysis for determining discrete partitions of cells or clusters, the Leiden algorithm [22] and K-Means [46]. Both methods have a hyperparameter to control the resolution or number of learned clusters, the resolution hyperparameter (res) for Leiden and K for K-Means. We assessed the results of these approaches on the simulated and real datasets described above, after normalizing and scaling the counts as is standard procedure for scRNAseq analysis [39]. For each count matrix, rows or cells were scaled to the same total count (10^4^) and log-transformed (log1p). We ran the Leiden or K-Means algorithms on the **U** and **S** matrices alone, as well as **U** + **S** (summed) matrices given standard practice in the 10x Genomics Cell Ranger 7.1.0 pipeline to sum intron- and exon-aligning reads, and **U** *⊕* **S** (concatenated) matrices which mimic the independent modeling of the two modalities by common integration methods [47]. The germ cell dataset (Fig. 1) was first reduced to 40 dimensions by PCA prior to clustering, following standard practice [45]. Leiden was run by default at res = 1.0 (as in scanpy), but results were additionally shown for a range of res and K values.

### Model Assessment and Metrics

To assess meK-Means inference over the epochs, we report the Q function (see Equation (4)) and Kullback-Leibler Divergence (KLD) (see Equation (2)), summed across each cluster k, at each epoch to determine convergence. KLDs are not directly comparable between models of various Ks.

#### Algorithm 1: meK-Means

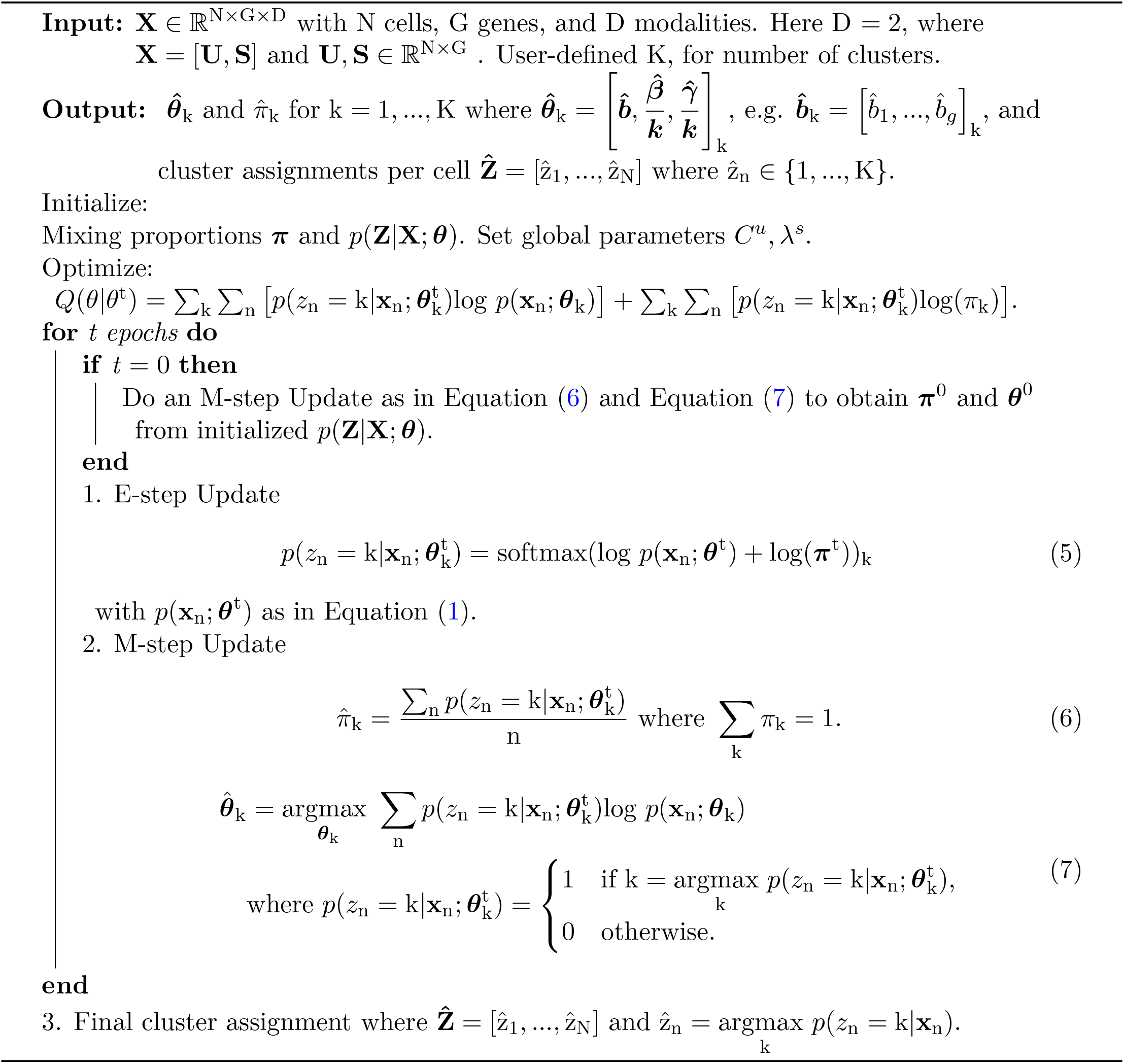

Standard error, *σ*, values for the inferred parameters are calculated from the square root of the diagonals of the inverse Fisher Information Matrix (FIM). The FIM is calculated as the inverse of the Hessian matrix of the KLDs between the observed count histograms and the distributions induced by the final inferred parameters. 99% confidence intervals (C.I.s) i.e. 2.756*σ*, are displayed for the relevant parameter plots in the figures.

We use three main criteria for rejecting genes which display poor model fits (possibly due to low counts and/or high variance). (1) Genes are rejected if inferred parameters remained close to the initialized parameter bounds prior to inference i.e. where gradient descent performance was poor and/or reliable estimates weren’t learned (as in [37]). These genes were not included in the generated figures. (2) We use the p-values obtained from Chi-squared tests between observed count histograms and the distributions induced by the inferred parameters (after Bonferroni correction for the number of genes tested), in conjunction with (3) Hellinger distances between these two distributions, to reject gene parameter fits with p-values *<* 0.05*/*G and Hellinger distances *>* 0.05 [37].

The Akaike Information Criterion (AIC) for model selection is calculated at the final epoch for all models, and is used to compare model fits over a range of K values (from meK-Means inference runs), defined as :

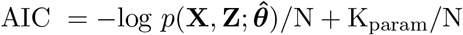

where K_param_ = 3 *×* G *×* K_Final_ + (K_Final_ *−* 1), for G genes and N cells, and where K_Final_ is the final number of clusters k that are assigned to cells. This metric thus incorporates general model fit (through the likelihood of the data given the parameters) and a penalty for the complexity of the model (through the number of parameters).

For determining differentially expressed genes at the parameter-level, DE-*θ* genes, log_2_FCs (log_2_ fold changes) are calculated for each gene between each parameter, given two clusters for comparison. ‘Markers’ denote DE-*θ* genes where log_2_FC *>* 2.0 (or *>* 1.3 for monocyte clusters comparisons).

For assessing the cluster assignments of the standard approaches, whether in comparison to ground truth clusters or between the results of the standard approaches for different input options, the Adjusted Rand Index (ARI) was used as a similarity measure between two cluster assignments. The ARI adjusts for expected similarity between random assignments, such that 1.0 denotes overlapping assignments (up to a permutation), and 0.0 denotes essentially random assignments (though negative values are possible for especially discordant assignments).

## Data and Code Availability

Data used in this study is available on [CaltechData](https://data.caltech.edu/search?q=mek-means&l=list&p=1&s=10&sort=bestmatc), with download links provided in Supplementary Table 1.

All code used to generate the figures and results in the paper, as well as a Google Colaboratory notebook with example usage of meK-Means, is available at https://github.com/pachterlab/CGP_2023. meK-Means is incorporated as a part of the *Monod* package [37] for single-cell, CME-based parameter inference, whose general documentation can be found here https://monod-examples.readthedocs.io/en/latest/.

## Acknowledgements

We thank Meichen Fang, Catherine Felce, and Laura Luebbert for helpful feedback on the manuscript and visualizations, and Ángel GÁlvez-MerchÁn for feedback on PBMC data gene selection. The meK-Means project was funded, in part, by NIH 5UM1HG012077-02.

## Notes

### Competing Interest Statement

The authors have declared no competing interest.

### Summary of Updates

Cell type names and gene symbol formats updated, to match convention

