## Supplementary Materials for "Biophysically Interpretable Inference of Cell Types from Multimodal Sequencing Data"

Tara Chari[1], Gennady Gorin[1], and Lior Pachter[1,2]

[1]Division of Biology and Biological Engineering,
California Institute of Technology, Pasadena, California

[2]Department of Computing and Mathematical Sciences,
California Institute of Technology, Pasadena, California

September 18, 2023

### Tables

#### Datasets

All datasets used in this study are listed in Supplementary Table 1, with links to the original data and other metadata/processed versions as necessary.

| Dataset | Technology | Full Data Download | Processed Data Download | Metadata Link |
|---|---|---|---|---|
| Testicular Germ Cell Data (E13) | 10xv2 | https://www.ncbi.nlm.nih.gov/geo/query/acc.cgi?acc=GSE136220 | https://doi.org/10.22002/e6mz6-r3g90 | https://doi.org/10.22002/b8xj7-0t796 |
| Mouse Primary Motor Cortex (MOp) | 10xv3 | https://zenodo.org/record/7497222/files/allen_B08_raw.loom?download=1 | https://doi.org/10.22002/zq7kd-57n29 | N/A (In processed loom file) |
| 10x Genomics 10k Human PBMCs | 10xv3 | https://zenodo.org/record/7388133/files/GP_2021_3_raw_data.tar.gz?download=1 | N/A | https://doi.org/10.22002/7fbwg-rx843 |

**Supplementary Table 1. Dataset Metadata.** Datasets used for all analyses. 'Processed' data refers to the gene-filtered or loom file form of the data, if the loom file was not already provided.

### Figures

#### Standard Clustering Demonstration

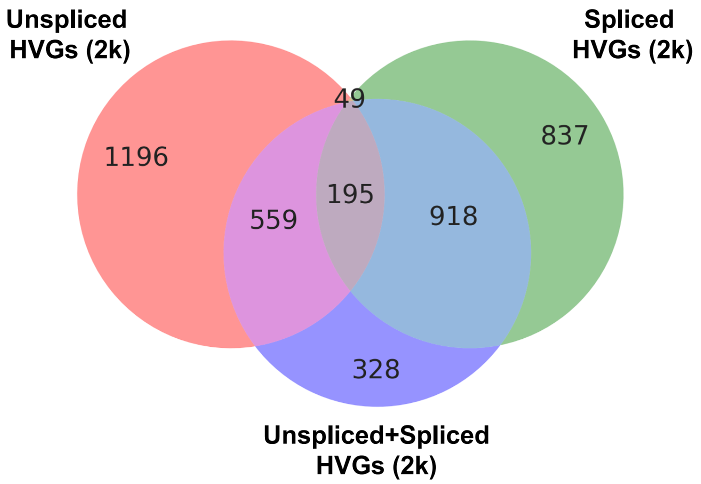

**Supplementary Fig 1. Overlap of Highly Variable Genes Between Input Options.** Venn diagram showing overlap of highly variable genes (HVGs), following standard selection of the top 2000 HVGs for each input matrix (given Unspliced and Spliced matrices) (see Methods).

### Simulated Data Analysis

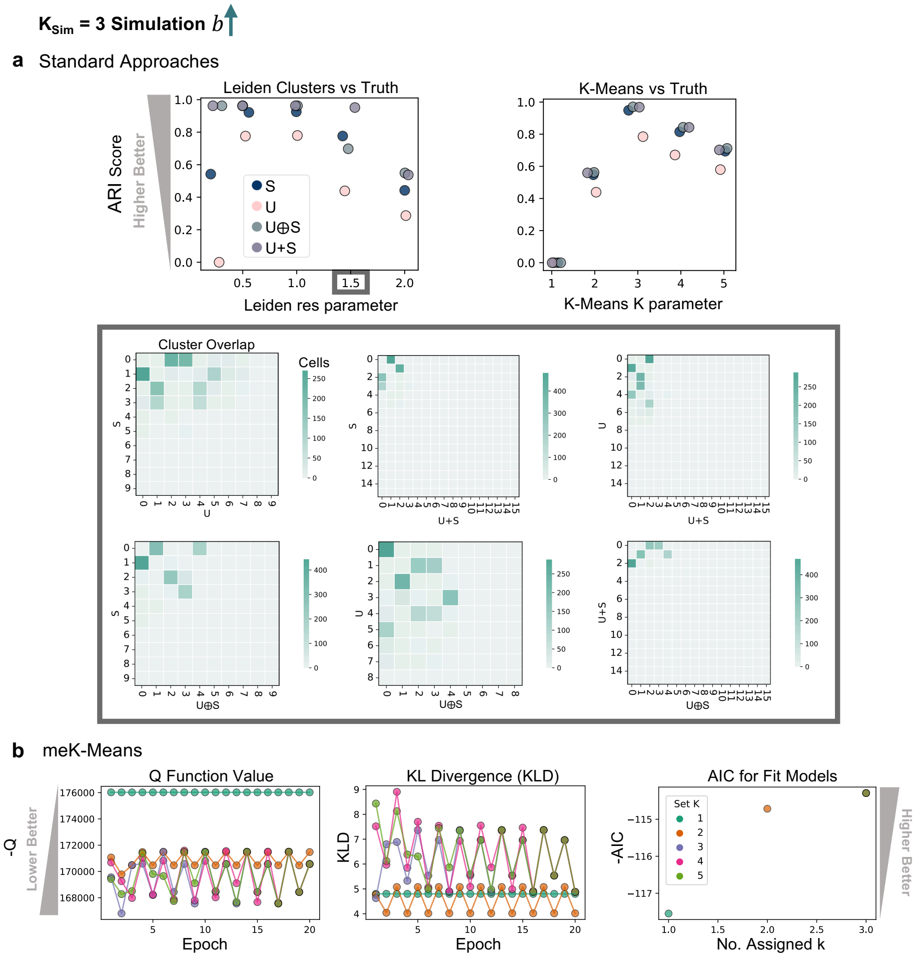

**Supplementary Fig 2. Burst Size Simulation Performance for Standard and meK-Means Approaches. a)** Top row, ARI scores for correspondence of learned clusters to the true clusters for $K_{Sim} = 3$ simulated data. Leiden results shown over possible res values, and K-Means results shown over possible K values. Grey box below shows confusion matrix/overlap between the results across the possible input options, for Leiden at res = 1.5. **b)** meK-Means inference results over possible values of K for the $K_{Sim} = 3$ simulated data. (Left to Right) Negative Q function (-Q) and Kullback-Leibler Divergence (KLD) over epochs, and negative Akaike Information Criterion (-AIC) versus the final k number of K clusters assigned (see Methods for metrics).

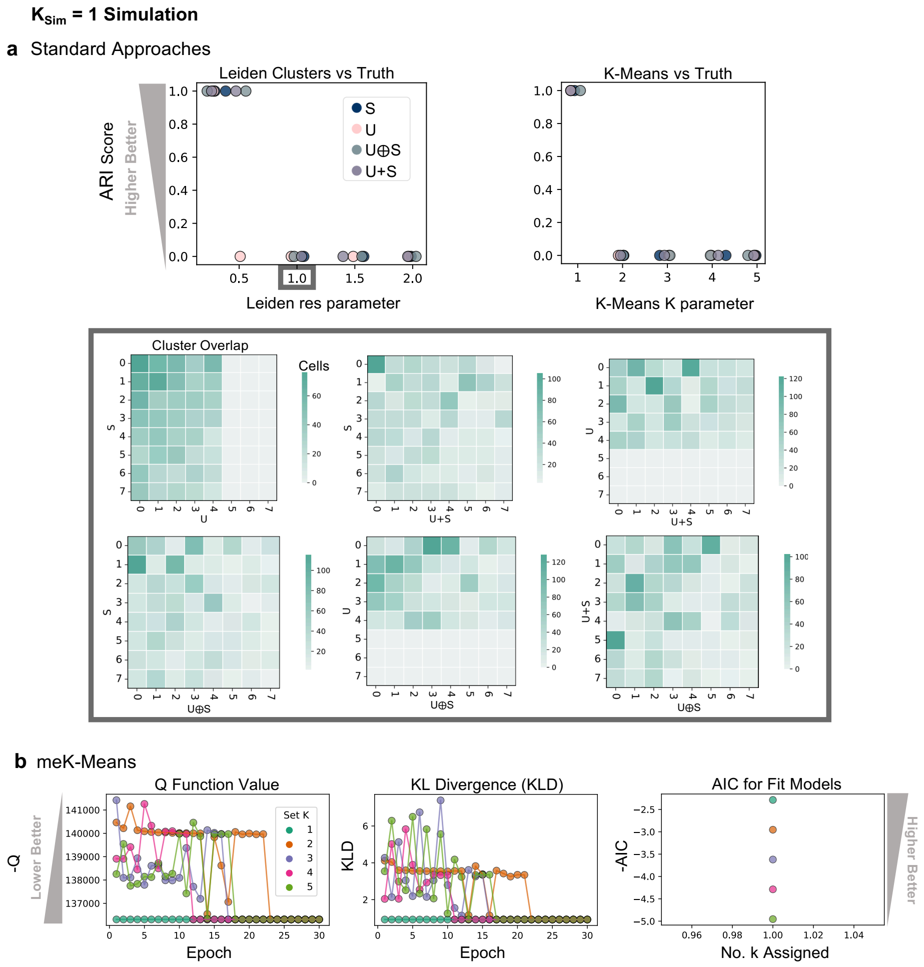

**Supplementary Fig 3. Negative Control Performance for Standard and meK-Means Approaches. a)** Top row, ARI scores for correspondence of learned clusters to the true clusters for $K_{\mathrm{Sim}} = 1$ simulated data. Leiden results shown over possible res values, and K-Means results shown over possible K values. Grey box below shows confusion matrix/overlap between results over the possible input options, for Leiden at res = 1.0 (default). **b)** meK-Means inference results over possible values of K for the $K_{\mathrm{Sim}} = 1$ simulated data. (Left to Right) Negative Q function (-Q) and Kullback-Leibler Divergence (KLD) over epochs, and negative Akaike Information Criterion (-AIC) versus the final k number of K clusters assigned (see Methods for metrics).

### MOp Data Analysis

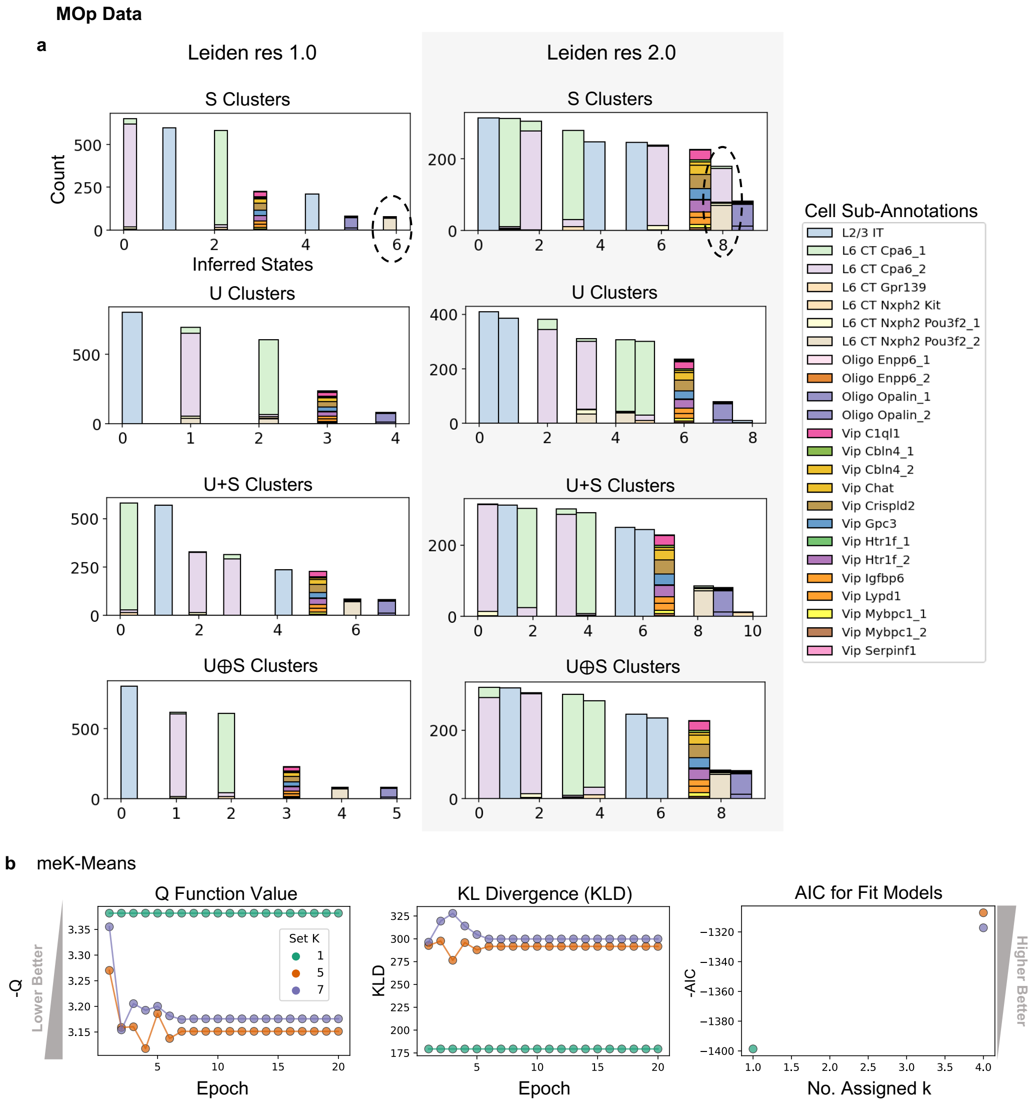

**Supplementary Fig 4. Performance of Standard and meK-Means Approaches on MOp Data. a)** Left column shows distributions of original, sub-type annotations across Leiden cluster assignments for each possible input (down the column), at res = 1.0. Right column shows same plots at Leiden res = 2.0. Example of changing assignments for the same annotated cells highlighted in dashed ovals. **b)** meK-Means inference results over possible values of K for the MOp data. (Left to Right) Negative Q function (-Q) and Kullback-Leibler Divergence (KLD) over epochs, and negative Akaike Information Criterion (-AIC) versus the final k number of K clusters assigned (see Methods for metrics).

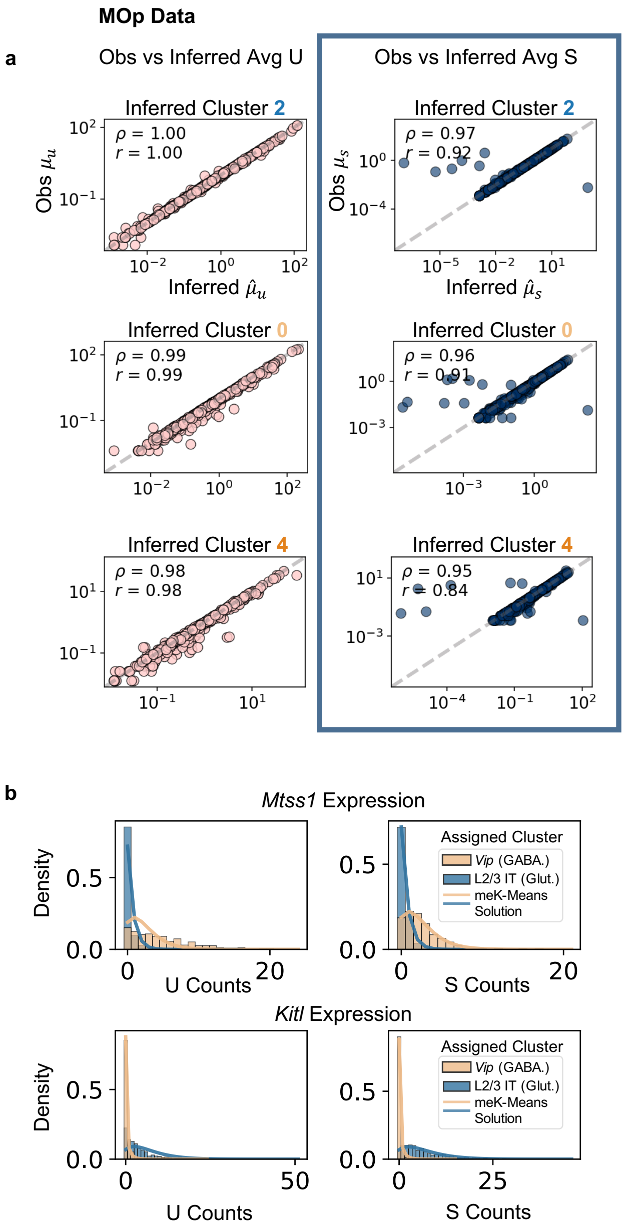

**Supplementary Fig 5. Correlation of meK-Means Parameters to Observed Means for MOp data. a)** Columns show Spearman ($\rho$) and Pearson ($r$) correlation of observed **U** (left) and **S** (right) means (e.g. $\mu_u$) versus the inferred count means (e.g. $\hat{\mu}_u$). Correlations shown for inferred clusters k $= 2, 0, 4$ down the columns. **b)** Marginal count distributions (histograms) for RBP-mediated genes in *Vip* and L2/3 IT clusters. Curves denote solution for the marginal distributions from the meK-Means parameters.

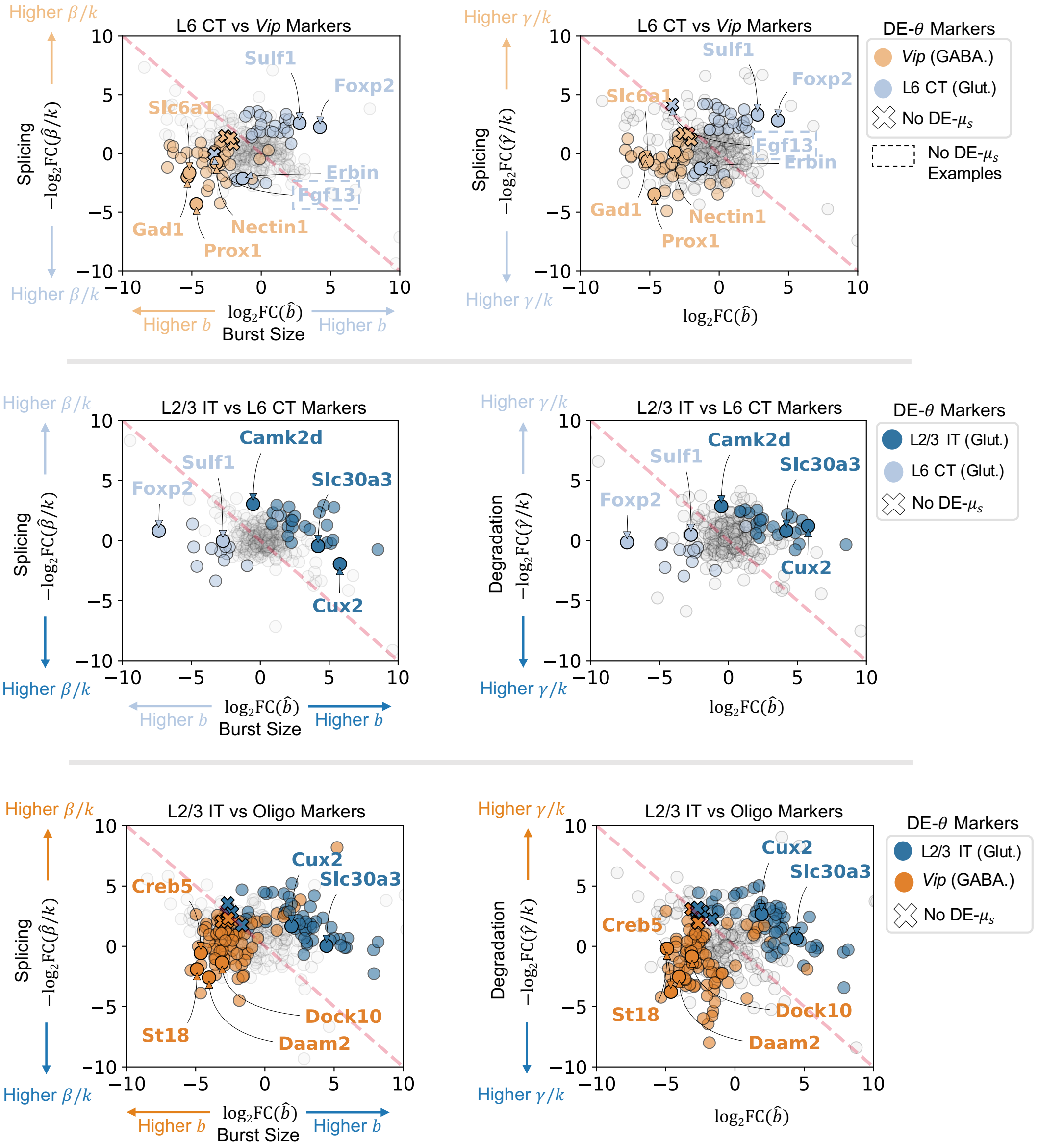

**Supplementary Fig 6. DE-$\theta$ Gene Markers for MOp Data.** DE-$\theta$ genes, based on $\log_2$ fold-changes (FC) between MOp clusters (see Methods). Left plots show splicing versus burst size FCs for denoted MOp cluster comparisons. Right plots show degradation versus burst size FCs for the same cluster comparisons. Genes denoted with Xs in the plots indicate parameter-level FCs not detected at the (observed) mean spliced-level (gene examples boxed in dashed lines)

**PBMC Data Analysis**

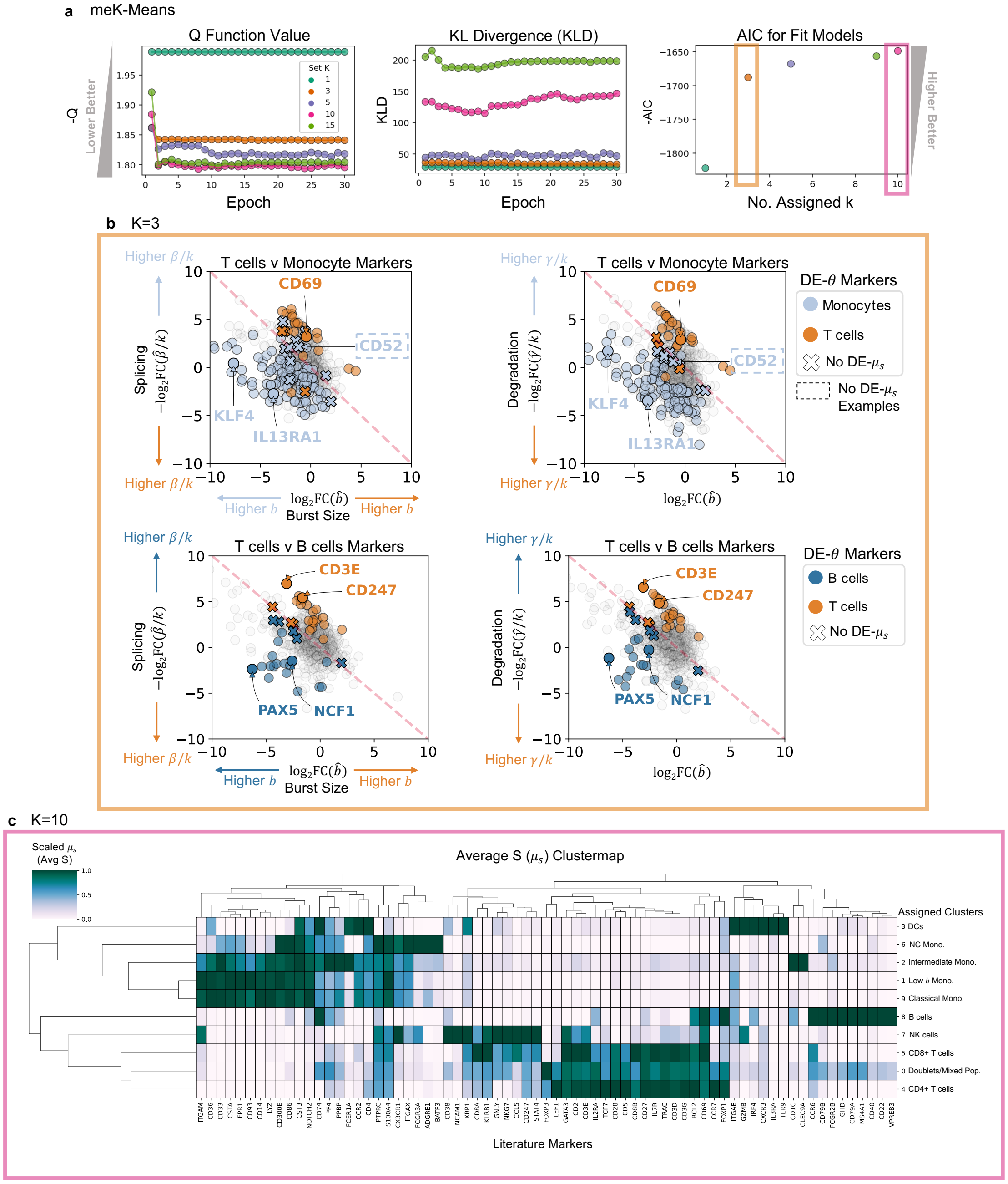

**Supplementary Fig 7. Initial Assignments of PBMC Data from meK-Means Results. a)** meK-Means inference results over possible values of K for the PBMC data. (Left to Right) Negative Q function (-Q) and Kullback-Leibler Divergence (KLD) over epochs, and negative Akaike Information Criterion (-AIC) versus the final k number of K clusters assigned (see Methods for metrics). **b)** DE-$\theta$ genes, based on $\log_2$ fold-changes (FC) between PBMC clusters, for meK-Means K=3. Left plots show splicing versus burst size FCs for denoted PBMC cluster comparisons. Right plots show degradation versus burst size FCs for the same cluster comparisons. Genes denoted with Xs in the plot indicate parameter-level FCs not detected at the (observed) mean spliced-level (gene examples boxed in dashed lines). **c)** Heatmap of inferred clusters from meK-Means with K=10, clustered by (observed) mean spliced expression of literature-based markers (not necessarily included in the meK-Means inference).

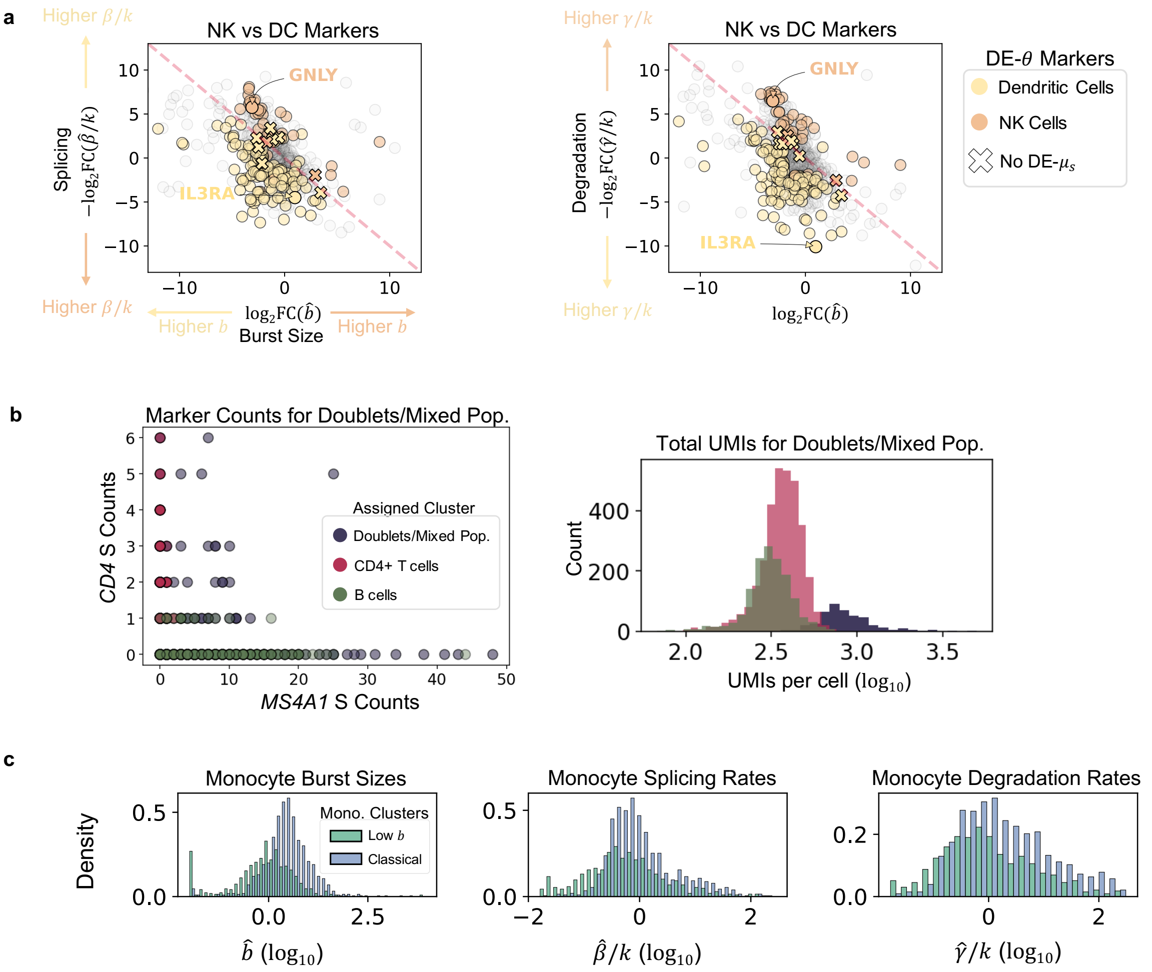

**Supplementary Fig 8. Analysis of meK-Means K=10 Results on PBMC Data. a)** DE-$\theta$ genes, based on $\log_2$ fold-changes (FC) between Natural Killer (NK) cells and Dendritic Cells (DCs), as denoted by the inferred clusters. Left plot shows splicing versus burst size FCs for the NK cells vs DC cells. Right plot shows degradation versus burst size FCs for the same cluster comparisons. Genes denoted with Xs in the plot indicate parameter-level FCs not detected between (observed) mean spliced counts. **b)** Observed **S** counts of known gene markers in the Doublet/Mixed Population alongside B and T cell expression of those markers. Right plot shows $\log_{10}$ total UMI counts per cell in each cluster. **c)** Distributions (histograms) of parameter values in $\log_{10}$ across genes shown for both the 'Low $b$ (burst size)' and 'Classical' monocyte clusters.

### Note

#### Properties of meK-Means Clustering

Here we describe the properties of meK-Means in relation to common clustering and mixture model-based methods, as meK-Means represents a unique clustering approach which utilizes various properties from these methodologies. In particular, the Expectation-Maximization (EM) approach in meK-Means is derived from a general mixture model (MM) formulation, but utilizes a more restricted variant akin to K-Means. K-Means is a classical method for partitioning observations into discrete states or clusters [1]. Here we use 'states' as a more general term to denote underlying, shared properties between observations. Where K-Means enforces 'hard' assignments of observations $\mathbf{X}$ (cells) into states, Gaussian MMs or GMMs (and MMs in general), can be viewed as probabilistic extensions which instead provide 'soft' assignment of observations to latent states $\mathbf{Z}$. The EM algorithm is commonly used to fit GMMs, iterating between updating posterior probabilities $p(\mathbf{Z}|\mathbf{X}, \boldsymbol{\theta}^{\mathrm{t}})$ (where $\boldsymbol{\theta}^{\mathrm{t}} = \boldsymbol{\mu}^{\mathrm{t}}, \boldsymbol{\Sigma}^{\mathrm{t}}$ are parameters of a *Normal* distribution) and updating $\boldsymbol{\theta} = \underset{\boldsymbol{\theta}}{\operatorname{argmax}} \sum_{\mathbf{Z}} p(\mathbf{Z}|\mathbf{X}, \boldsymbol{\theta}^{\mathrm{t}}) \log p(\mathbf{X}, \mathbf{Z}|\boldsymbol{\theta})$.

Notably, the K-Means algorithm can be interpreted as a limiting case of the EM algorithm for GMMs, where we treat the data as coming from a mixture of spherical Gaussians [2]. As the trace of the covariance matrix ($\boldsymbol{\Sigma}$) approaches zero, the posterior probabilities ($p(\mathbf{Z}|\mathbf{X}, \boldsymbol{\theta})$) converge on binary assignments of observations to states (where only one state has a probability of 1, and all others 0, for an observation) [2]. This binary, or hard, assignment thus distinguishes the K-Means algorithm, from the soft assignment of states in the standard GMM.

meK-Means is named as such ('mechanistic K-Means') as it utilizes a similar hard assignment approach in the M Step, for obtaining MLE parameter estimates. Though we define meK-Means as a mixture model over the full, joint distribution of observations $\mathbf{X}$ and states $\mathbf{Z}$ (using the CME formalism), binary assignment of cells is used during the M step (see Equation (7)) to delineate distinct, observed count histograms for each state for the numerical, CME parameter estimation procedure (see Equation (2)) . However, meK-Means does retain probabilistic interpretation of the learned states e.g. the ability to assess the likelihood of the observations $\mathbf{x}_{\mathrm{n}}$ with respect to the K states, and enables statistical analysis of the inferred models.

In terms of the assumptions underlying K-Means, GMMs, and meK-Means, both K-Means and GMMs assume a Gaussian representation of the data, though single-cell data is sparse, discrete count data (often modeled with Poisson or Negative Binomial distributions) [3]. MMs can be extended to such discrete count distributions, however the CME approach provides a natural way to *integrate* modalities, which is not inherent in such MM methods as modalities are often treated independently, and how 'best' to model them together is undefined. Additionally, with the CME formalism, we infer parameters which more explicitly represent biological processes, as opposed to descriptive parameters which represent the observed counts only (e.g. mean expression values) lacking a connection to underlying, generative processes.

Another key difference between meK-Means and clustering methods such as K-Means or (G)MMs, is the nature of the input and output. The input data $\mathbf{X}$ is a multidimensional array (or tensor) of cells-by-genes-by-modalities (i.e. $\mathbf{X} \in \mathbb{R}^{\mathrm{N \times G \times D}}$ with N cells, G genes, and D modalities as described in Methods). In this case, the features (modalities) are not vectorized or flattened to a matrix and converted to a distance or affinity matrix for partitioning as is the case with centroid-based clustering methods such as K-Means [4]. Rather, use of the full tensor allows for retention of structural properties/patterns between the dimensions or modalities [5]. This parallels other work in joint clustering of high-dimensional tensor data where the objective is to simultaneously learn cluster labels and estimate model parameters [5]. As described in [5], from a dataset $\mathbf{X} \in \mathbb{R}^{d_1 \times d_2 \times d_3}$ for K mixtures, one would learn mixture weights $\pi_k$, cluster assignments $\boldsymbol{z} = (z_1, ..., z_n)$ for $n$ tensor observations, and parameters of the governing multivariate Gaussian $\boldsymbol{U}_k, \boldsymbol{\Sigma}_k$ where $\boldsymbol{U}_k \in \mathbb{R}^{d_1 \times d_2 \times d_3}$ and $\boldsymbol{\Sigma}_k = \{\boldsymbol{\Sigma}_{k,1}, \boldsymbol{\Sigma}_{k,2}, \boldsymbol{\Sigma}_{k,3}\}$ and $\boldsymbol{\Sigma}_{k,1} \in \mathbb{R}^{d_1 \times d_1}$ [5]. However, unlike [5] the modalities in meK-Means are not modeled as multivariate Gaussian distributions. Rather, the CME formalism defines shared parameters over these modalities which in turn induce their joint distribution. Thus from an input of $\mathbf{X} \in \mathbb{R}^{\mathrm{N} \times \mathrm{G} \times \mathrm{D}}$, we effectively learn an output $\mathbf{O} \in \mathbb{R}^{\mathrm{N} \times \mathrm{G} \times \mathrm{P}}$, where P is the number of parameters in $\boldsymbol{\theta}$ (P=3 for parameters $b, \beta, \gamma$) and the $N$ cells are partitioned into k $\leq$ K layers (see Supplementary Note Fig. 1). Such a framework can in principle be extended to higher-dimensional tensors, where for example $\mathbf{X} \in \mathbb{R}^{\mathrm{N} \times \mathrm{C} \times \mathrm{G} \times \mathrm{D}}$ with C conditions e.g. environmental perturbations or drug treatments.

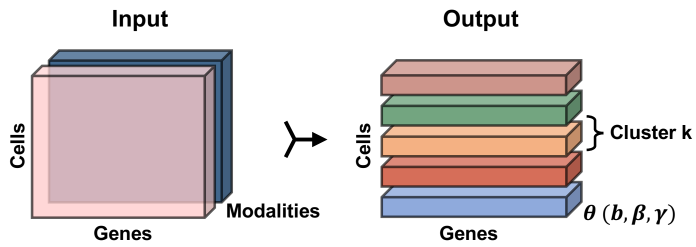

**Supplementary Note Fig 1. meK-Means for Joint Clustering of Data.** Diagram of tensor input for meK-Means, and the resulting partitions and parameters learned simultaneously from the data.
